# MicroRNA-27a is a key modulator of cholesterol biosynthesis

**DOI:** 10.1101/383448

**Authors:** Abrar A. Khan, Heena Agarwal, S. Santosh Reddy, Vikas Arige, Vinayak Gupta, Ananthamohan Kalyani, Manoj K. Barthwal, Nitish R. Mahapatra

## Abstract

Hypercholesterolemia is a strong predictor of cardiovascular diseases. 3-Hydroxy-3-methylglutaryl-coenzyme A reductase gene (*Hmgcr*) coding for the rate-limiting enzyme in the cholesterol biosynthesis pathway is a crucial regulator of plasma cholesterol levels. However, the post-transcriptional regulation of Hmgcr remains poorly understood. The main objective of this study was to explore the role of miRNAs in the regulation of *Hmgcr* expression. Systematic *in silico predictions* and experimental analyses reveal that miR-27a specifically interacts with the *Hmgcr* 3’-untranslated region in murine and human hepatocytes. Moreover, our data shows that *Hmgcr* expression is inversely correlated with miR-27a levels in various cultured cell lines, human and rodent tissues. Actinomycin D chase assays and relevant experiments demonstrate that miR-27a regulates Hmgcr by translational attenuation followed by mRNA degradation. Early Growth Response 1 (Egr1) regulates miR-27a expression under basal and cholesterol-modulated conditions. miR-27a augmentation via tail-vein injection of miR-27a mimic in high cholesterol diet-fed *Apoe*^−/−^ mice shows down-regulation of hepatic Hmgcr and plasma cholesterol levels. Pathway and gene expression analyses show that miR-27a also targets several other genes (apart from *Hmgcr*) in cholesterol biosynthesis pathway. Taken together, miR-27a emerges as a key regulator of cholesterol biosynthesis and has therapeutic potential for clinical management of hypercholesterolemia.

## INTRODUCTION

Cardiovascular diseases (CVDs) remain the leading cause of global mortality and morbidity [1]. Among various determinants of CVDs, plasma cholesterol is an important factor contributing to multiple disease states including atherosclerosis, coronary artery disease, obesity, hypertension and type 2 diabetes [2–4]. 3-hydroxy-3-methylglutaryl-coenzyme A (HMG-CoA) reductase gene (human: *HMGCR*, mouse/rat: *Hmgcr*) that codes for a ~ 97 kDa endoplasmic-reticulum membrane glycoprotein catalyzing the rate-limiting step in the cholesterol biosynthesis pathway [5] is, therefore, a critical modulator of dyslipidemia and consequent CVDs.

Statins, HMG-CoA reductase inhibitors, are widely used to reduce high cholesterol levels and risk of CVDs [6]. HMGCR expression/enzyme activity is modulated by feedback control mechanisms involving sterols and non-sterols at transcriptional and post-translational levels by family of sterol regulatory element binding proteins (SREBPs), SREBP cleavage activated protein (SCAP), and insulin induced genes (Insig1 and Insig2) [7]. However, the molecular mechanisms regulating *Hmgcr* expression at the post-transcriptional level are poorly understood.

MicroRNAs (miRNAs) are small noncoding RNAs that control gene expression by translational repression and/or mRNA degradation. Recent studies have revealed that miRNAs play important roles in cardiovascular physiology, and pathophysiology [8,9]. We performed systematic computational and extensive experimental analyses that revealed a crucial role for miR-27a in the post-transcriptional regulation of *Hmgcr* under basal and elevated cholesterol conditions. This study also highlights the previously unknown role of Egr1 in miR-27a expression. Interestingly, miR-27a targets multiple genes in addition to *Hmgcr* in cholesterol biosynthesis. In line with these *in vitro* findings, miRNA-27a represses Hmgcr expression in liver tissues and modulates plasma cholesterol levels in high cholesterol diet-fed *Apoe* ^−/−^ mice.

## MATERIALS AND METHODS

### Comparative genomics analyses

For rat QTL analysis, data pertaining to elevated lipid/cholesterol-QTLs, their respective LOD scores and all the genes with their respective positions in a particular QTL was mined from the Rat Genome Database (Table S1). Each QTL is designated a ‘Logarithm of odds’ (LOD) score which is a measure of strength of evidence for the presence of a QTL at a particular location and its association to a trait. The elevated lipid/cholesterol-QTLs in rats were plotted against the respective LOD scores. We also compared mouse and rat *Hmgcr* gene sequences (GenBank accession no: NM_008255 and NM_013134.2, respectively) using mVISTA browser.

### *In silico* predictions of potential miRNA binding sites in *Hmgcr*-3′UTR and putative miRNA targets in the cholesterol biosynthesis pathway

Putative miRNA binding sites in mouse Hmgcr (*Hmgcr*)-3’UTR sequence (NCBI reference number: NM_008255.2) were predicted using various bioinformatic algorithms [viz. miRWalk, miRanda, TargetScan, PITA, RNA22 and RNAhybrid (Table S2)]. Since, a large number of miRNAs were predicted by these online tools, we selected only those miRNAs that were predicted by at least five algorithms. Further, differences in hybridization free energy indicating the stability of the microRNA-mRNA interaction was determined computationally by two online tools called PITA and RNAhybrid. The lower or more negative ΔΔG value predicted by PITA indicates stronger binding of the microRNA to the given site; as a rule of thumb, sites having ΔΔG values below −10 are likely to be functional and selected for experimental validation. RNAhybrid calculates the minimum free energy (ΔG) of hybridization between target mRNA and miRNA. An RNAhybrid ΔG score of less than −20 kcal/mol is a strong indicator for interactions of a miRNA with target mRNA. In each case, the minimum number of nucleotides in seed sequence was selected as 6 and no mismatch or G:U wobble base-pairing was allowed in the region of seed sequence. The lower or more negative ΔG (<−10) or ΔΔG (<−20) values predicted by PITA/RNAhybrid, in general, indicate a stronger binding of the miRNA to the target gene 3’-UTR.

In order to predict miR-27a target genes in the cholesterol biosynthesis pathway, putative miR-27a targets were retrieved from TargetScan and miRWalk (Table S2). These targets were grouped by PANTHER classification system (http://www.pantherdb.org/) based on their molecular functions and the gene IDs mapping to the cholesterol biosynthesis pathway were selected. In addition, genes involved in the cholesterol biosynthesis pathway having an RNAhybrid ΔΔG score of less than −20 were also considered as putative miR-27a targets.

### Tissue-specific expression of endogenous *HMGCR*, hsa-miR-27a-3p, hsa-miR-28 and hsa-miR-708 in human tissues

For correlation analysis, human tissues that showed consistent *HMGCR* expression across GTEx, BioGPS and SAGE (Table S2) databases were selected for analysis. The expression of *HMGCR* across various human tissues was mined from the GTEx portal (Table S2). Likewise, tissue-specific hsa-miR-27a-3p expression was obtained from DASHR (Table S2). Further, tissues showing hsa-miR-27a-3p expression in the range of 100-1000 RPM were selected since miRNAs expressed in this range were reported to bring about significant target repression [10]. Only tissues common to the GTEx portal and DASHR were chosen for correlation analysis. The data was normalized to a particular tissue (having the lowest expression of hsa-miR-27a-3p) and expressed as fold change. Specifically, the expression data from DASHR and GTEx was normalized to spleen. Likewise, the tissues that were used for correlation analysis of hsa-miR-27a-3p and *HMGCR* expression were also analyzed for hsa-miR-28 and hsa-miR-708 expression.

### Pathway analysis for miR-27a targets

In order to identify pathways that may be targeted by miR-27a, TargetScan and mirPath v3. tools were employed (Table S2). TargetScan relies on an *in silico* prediction of miRNA targets in a specific pathway while mirPath v3. employs prediction algorithms and experimentally validated miRNA-gene interactions. Interestingly, 21 pathways that were predicted miR-27a targets were commonly enriched in both these tools including steroid biosynthesis, fatty acid metabolism and biosynthesis.

### Generation of *Hmgcr* 3’-UTR/luciferase, mmu-miR-27a promoter/luciferase reporter constructs and miRNA expression plasmids

To generate the *Hmgcr* 3’-UTR/luciferase construct, mouse *Hmgcr* 3’-UTR domain (+20359/+21975 bp) was PCR-amplified using Phusion® High Fidelity DNA Polymerase (Finnzymes), mouse genomic DNA (Jackson Laboratory, Bar Harbor, USA) and gene specific primers (FP: 5’-CGT**<u>GCTAGC</u>**GGATCCTGACACTGAACTG-3’, RP: 5’-GC**<u>GGCCGGCC</u>**TTCAATGTTAACTTCCTTTC -3’). The numberings of the nucleotide positions are with respect to cap site as +1 position. Bold and underlined nucleotides in forward and reverse primers are the restriction sites for *Nhe*I and *Fse*I, respectively, that were added to enable cloning of the PCR-amplified 3’-UTR into the firefly-luciferase expressing pGL3-promoter reporter vector (Promega). The purified *Hmgcr* 3’-UTR PCR product was cloned between *Xba*I and *Fse*I sites of pGL3-Promoter vector because *Nhe*I digested PCR product had compatible ends for *Xba*I-digested pGL3-promoter reporter vector. Authenticity of the Hmgcr 3’-UTR reporter plasmid was confirmed by DNA sequencing using pGL3-promoter vector sequencing primers [forward primer (1782-1801bp): 5′-CGTCGCCAGTCAAGTAACAA-3′ and reverse primer (2118–2137bp): 5′-CCCCCTGAACCTGAAACATA-3′)]; the resultant plasmid was named as m*Hmgcr* 3’UTR.

To abrogate the binding of miR-27a, 3’UTR-deletion construct was generated using site-directed mutagenesis wherein the putative miR-27a binding site was deleted. The 3’UTR-deletion construct for miR-27a was generated by using the wild-type Hmgcr-3’UTR-reporter construct as template and the following primers: forward, 5’-CGCGGGCATTGGGTTCTCAATTAAAAATCTCAATGCACT-3’ and reverse, 5’-AGTGCATTGAGATTTTTAATTGAGAACCCAATGCCCGCG-3’. This deletion in the reporter plasmid was confirmed by DNA sequencing and the resultant construct was named as m*Hmgcr* 27a mut 3’UTR.

The mmu-miR-27a promoter/luciferase reporter construct was generated by amplifying the −1079 bp to +26 bp region of the miR-27a promoter using mouse genomic DNA as described above and primers (forward, 5’-CTA**<u>GCTAGC</u>**AACTTTAACTGGCACGCAGG-3’, and reverse, 5’-CCG**<u>CTCGAG</u>**GGCATCAAATCCCATCCC-3’). Bold and underlined nucleotides in forward and reverse primers are the restriction sites for *Nhe*I and *Xho*I, respectively, that were added to assist in cloning of the PCR-amplified miR-27a promoter region into the pGL3-basic vector (Promega). The authenticity of resultant construct (referred to as miR-27a promoter) was confirmed by DNA sequencing.

To generate plasmid expressing miR-27a, the sequence of the pre-miRNA was retrieved from miR-Base/UCSC genome browser, PCR-amplified using mouse genomic DNA as template and primers [forward, 5’-CGC**<u>GGATCC</u>**TCGCCAAGGATGTCTGTCTT-3’ and reverse, 5’-CCG**<u>CTCGAG</u>**GTTTCAGCTCAGTAGGCACG-3’]. Bold and underlined nucleotides are *BamH*I and *Xho*I restriction sites that were added to the forward and reverse primers, respectively. The purified PCR-amplified DNA was cloned between *BamH*I and *Xho*I restriction sites in pcDNA3.1 vector (Invitrogen). The resultant plasmid was named as miR-27a expression plasmid and the accuracy of the insert was confirmed by DNA sequencing using miRNA specific primers. Similarly, miR-27b and miR-764 expression plasmids were generated using specific primers [forward, 5’-CGC**<u>GGATCC</u>**TGCAGTTTGGAGAACAGAGG-3’ and reverse, 5’-CCG**<u>CTCGAG</u>**CTTGAGGCAGGCTGGTCT-3’; forward, 5’-CGC**GGATCC**CCTTGTGGTATTGTTGGAAGG-3’ and reverse, 5’-CCG**<u>CTCGAG</u>**TCTTTCCTTTGCTCTACCTTG-3’], respectively, harboring *BamH*I and *Xho*I restriction sites to enable cloning in pcDNA3.1 vector (Invitrogen).

### Cell lines, transfections and reporter assays

AML12 (alpha mouse liver 12; a gift from Dr. Rakesh K. Tyagi, Jawaharlal Nehru University, New Delhi, India), mouse neuroblastoma N2a and human hepatocellular carcinoma HuH-7 cells (obtained from the National Center for Cell Sciences, Pune, India) were cultured in Dulbecco’s modified Eagle’s medium with high glucose and Glutamine (HyClone), supplemented with 10% fetal bovine serum (Invitrogen, USA), penicillin G (100 units/mL), and streptomycin sulfate (100 mg/mL) (Invitrogen) in 25 cm^2^ tissue culture flasks (NEST) at 37 °C with 5% CO_2_. These cell lines were tested for mycoplasma infection routinely and treated with BM-Cyclin (Merck) to eliminate mycoplasma, if detected.

AML12 and HuH-7 cells were grown up to 70% confluence in 12-well plates and co-transfected with different doses (125, 250 and 500 ng) of miR-27a expression plasmid, along with 500 ng/well of m*Hmgcr* 3’UTR plasmid by Lipofectamine 2000 (Invitrogen) according to manufacturer’s instructions. Similarly, m*Hmgcr* 3’UTR plasmid or m*Hmgcr* 27a mut 3’UTR construct was co-transfected with miR-27a expression plasmid in a dose-dependent manner. In all these co-transfection experiments, the insert-free vector pcDNA3.1 was used as balancing plasmid. After 4 hrs of transfection, the culture media was changed with fresh complete media.

In other co-transfection experiments, AML12 cells were transfected with different doses of Egr1 expression plasmid (obtained from Dr. Wong, [11]) or Egr1 shRNA expression plasmids (obtained from Dr. Xiao, [12]) along with 500 ng/well of the miR-27a pro construct. In order to maintain equal amount of DNA across transfections, pcDNA3.1/pU6 was used as a balancing plasmid with Egr1/Egr1 shRNA expression plasmid, respectively.

In all co-transfection experiments, cells were lysed 36 hrs post-transfection and cell lysates were assayed for luciferase activity. Luciferase assays were carried out as described previously [13,14]. Total protein per individual well was also estimated in the same cell lysate using Bradford reagent (Bio-Rad). The reporter activities were normalized with the total protein content and expressed as luciferase activity/μg of protein or % over control.

### Animals and tissue samples

All animal-related procedures were approved by the Institutional Animal Ethics Committee at Indian Institute of Technology Madras as well as CSIR-Central Drug Research Institute, Lucknow and performed in accordance with the *Guide for the care and use of laboratory animals* published by the US National Institutes of Health (eighth edition) and the ARRIVE (Animal in Research: Reporting In Vivo Experiments) guidelines [15]. Male *Apoe* ^−/−^ mice (18–20 g; 8–10 weeks months old) on a C57BL/6 background were obtained from National Laboratory Animal Centre, CSIR-Central Drug Research Institute, Lucknow India. The animals were maintained in polypropylene cages under controlled room temperature at 25 °C on a 12 hr light-dark cycle. The mice were fed a high cholesterol diet (HCD) containing 0.21%, 20%, 50% and 21% cholesterol, protein, carbohydrate and fat (by weight), respectively (Research Diets) for 10 weeks. All mice received water and food *ad libitum.* After completion of 10 weeks of diet regimen, the mice were randomly assigned to three groups: HCD-fed mice injected with either saline (n=7) or miRVANA miR-27a mimic (Invitrogen; n=7) or miRVANA negative control oligos (Invitrogen; n=7), respectively.

The miR-27a mimic and negative control oligos were synthesized from Invitrogen, complexed with Invivofectamine 3.0 (Invitrogen) and injected via tail vein at a dose of 5 mg/kg body weight twice over a period of 11 days (second injection after a week of the first dosing). Invivofectamine 3.0 is a lipid nanoparticle delivery system used for the *in vivo* delivery of miRNAs/siRNAs in rodents. Owing to its safety and efficacy, multiple studies have reported to use of Invivofectamine 3.0 for delivery of miRNA mimic/inhibitors *in vivo* in mice [16–19]. For the *in vivo* delivery of the miR-27a mimic/control oligo, 75 μl of 19.2 μg/μl solution of control oligo/miR-27a mimic was added to equal volume of complexation buffer. This mixture was then diluted with equal volume of Invivofectamine 3.0 and vortexed for a few seconds to ensure miR-27a-Invivofectamine complex formation. The complexes were incubated at 50 °C for 30 min. These complexes were then diluted appropriately with PBS to attain a final concentration of 0.5 mg/ml. Around 200 μl of this formulation was injected into *Apoe* ^−/−^ mice via tail-vein. The injected mice were still continued on a high cholesterol diet (HCD) until they were sacrificed. The body weight of the animals was monitored every alternate day. After 4 days of the second dosing, the animals were euthanized by CO2 inhalation followed by cardiac puncture (for blood collection into heparin vials) and the organs were collected in RNAlater (Thermo Fisher) and neutral buffered saline and stored in −80 °C for further analyses. Wistar female rats at the age of 6 weeks were obtained from the King Institute of Preventive Medicine (Chennai, India). Liver, kidneys, heart and skeletal muscle were isolated following standard procedures.

### Analysis of biochemical parameters in plasma, blood and tissue samples from the miRNA/ control oligo-treated mice

Animals were fasted overnight and plasma was collected by spinning the whole blood at 2000 rpm at 4°C for 20 min for measuring total, HDL, LDL cholesterol levels using commercial kits (Randox) as per manufacturer’s instructions. Plasma ALT (alanine aminotransferase), AST (aspartate aminotransferase) and CK (creatine kinase) were also measured using kits (Randox) according to the manufacturer’s protocol. For estimating hepatic cholesterol levels, liver tissue (~20 mg) was homogenized in a solvent containing hexane: isopropanol in the ratio 3:2. The lipids were then extracted and tissue total cholesterol was assayed by Amplex Red cholesterol assay kit (Thermo Fisher) as per manufacturer’s protocol. For fasting blood glucose measurements, blood was collected via retro-orbital bleed after fasting the animals overnight and the blood glucose levels were measured using an Accu-Chek Active Blood Glucometer (Roche).

### RNA extraction and Real-time PCR

Total RNA was extracted from cell lines and tissue samples by using TRIzol (Invitrogen) as per the manufacturer’s instructions. cDNA synthesis was performed using High Capacity cDNA Reverse Transcription Kit (Applied Biosystems) and miR-27a/miR-27b/U6-specific stem-loop (SL; Table S3) or random hexamer primers. Quantitative real-time PCR (qPCR) was carried out using the DyNAmo HS-SYBR Green qPCR Kit (Finnzymes) and miR-27a/b primers and a universal reverse primer or gene specific primers (Table S3). The same mmu-miR-27-forward primer was also used to probe for miR-27b expression.

In certain experiments, AML12 cells were transfected with different doses of miR-27a expression plasmid or 1 μg of miRVANA miR-27a mimic (Invitrogen) and miRVANA negative control oligos (Invitrogen) or 60 nM of locked nucleic acid inhibitor of 27a (LNA 27a) and negative control oligos (Exiqon) using lipofectamine. Over-expression or down-regulation of miR-27a was determined by qPCR analysis.

For cholesterol depletion, AML12 cells (grown in 12 well plates) were treated with increasing doses (0, 1, 3.6 and 5 mM) of cholesterol-depleting reagent methyl-β-cyclodextrin (HiMedia) for 15 min. Cholesterol depletion was carried out in serum-free DMEM medium. Following cholesterol depletion, media was changed with fresh serum-free media and the cells were incubated for 6-9 hrs at 37°C in CO_2_ incubator. In another series of experiments, AML12 cells were treated with exogenous cholesterol (20 μg/ml) in serum free media for 6-9 hrs. Next, the cells were processed for RNA isolation followed by qPCR to measure the relative abundance of miR-27a and *Hmgcr* transcript. Total miRNAs isolated from the treated and control cells were subjected to cDNA synthesis followed by qPCR analysis probing for miR-27a and U6 RNA using miR-27a and U6 specific primers. In all the qPCR analysis, the relative abundance of miR-27a and *Hmgcr* was determined by calculating 2^−ΔΔCt^ of each reaction [20].

### Filipin staining of hepatocyte cells

AML12 cells were seeded at 60-70% confluency in 12/ 24 well plates overnight at 37 °C with 5% CO_2_. The cells were then treated with 20 μg/ml of exogenous cholesterol or 5mM of cholesterol-depleting reagent methyl-β-cyclodextrin (MCD) for 6 hrs or 15 min, respectively. Post-MCD treatment, the media was changed to serum-free media and the cells were incubated at 37 °C for 6-9 hrs. Following treatment with cholesterol or MCD, the cells were washed with PBS and fixed with 3.6% formaldehyde in PBS for 10 min at room temperature. The fixed cells were washed with PBS and stained with 50 μg/ml of Filipin III (a fluorescent dye that binds to free cholesterol) in the dark at room temperature for 2 hrs. Following fixing, the cells were washed with PBS again and imaged using an Olympus U-RFL-T fluorescence microscope (Olympus).

### mRNA stability assays

Actinomycin D, an extensively used and highly specific transcriptional inhibitor, was used to determine the Hmgcr mRNA stability as described earlier [21]. In this experiment, AML12 cells were transfected with miR-27a plasmid or pcDNA3.1. After 12 hrs of transfection, cells were treated with actinomycin D (5 μg/ml) (Himedia) for different time points (0, 6, 12 and 24 hrs). In both the cases (with/without miR-27a over-expression), Hmgcr mRNA decay was monitored by measuring the *Hmgcr* levels by qPCR. Hmgcr mRNA half-life was determined by using t_1/2_= −t (ln2)/ln(N_t_/N_0_) where N_t_= mRNA remaining at a specific time t, and N_0_= mRNA abundance in the beginning. Western blot analysis for Hmgcr level at each time point following Actinomycin D treatment was also performed.

### Ago2-Ribonucleoprotein immunoprecipitation (RIP) assays

Ago2-RIP assays were performed as described previously [22]. HuH-7 cells grown at 60% confluence in 100 mm dishes were transfected with 5 μg of miR-27a plasmid or pcDNA3 using Targetfect F2 transfection reagent (Targeting Systems). After 24 hrs of transfection, cells were lysed in 100 μl of ice cold polysome lysis buffer [5 mM MgCl_2_, 100 mM KCl, 10 mM HEPES (pH 7.0) and 0.5% Nonidet P-40] with freshly added 1 mM DTT, 100 U/ml recombinant ribonuclease (Takara) supplemented with protease inhibitor cocktail (Sigma-Aldrich) by tapping every 5 min for 3 secs over a period of 15min on ice. The lysates were then centrifuged at 14,000 rpm at 4 °C for 10 min. Supernatant was mixed with 900 μl of ice-cold NT2 buffer [50 mM Tris (pH 7.4), 150 mM NaCl, 1 mM MgCl_2_, 0.05% Nonidet P-40] containing freshly added 200 U/ml recombinant ribonuclease (Takara), 1 mM DTT, 15 mM EDTA. The lysates were pre-cleared with Rec protein G-Sepharose 4B beads (Invitrogen). Pre-cleared samples were then immunoprecipitated by overnight incubation at 4 °C with either 0.75 μg of anti-Ago2 antibody (abcam, ab57113) or non-immune mouse IgG (Sigma, I5831). On the following day, beads were washed five times with ice-cold NT2 buffer and divided into two fractions-one for RNA isolation to identify miRNA target genes and another for Western blotting to check for successful immunoprecipitation of Ago2. Anti-Ago2 antibody at a dilution of 1:2500 was used for Western blot analysis. RNA was isolated from the other fraction of beads by TRIzol (Invitrogen) followed by purification via Nucleospin miRNA columns (Machery-Nagel).

### Western Blot Analysis

AML12 or HuH-7 cells after transfection/cholesterol treatment/ depletion experiments were lysed in radioimmunoprecipitation assay (RIPA) buffer [50 mM Tris–HCl (pH 7.2), 150 mM NaCl, 1% (v/v) Triton X-100, 1% (w/v) sodium deoxycholate, 1mM EDTA and 0.1% (w/v) SDS] supplemented with 1mM PMSF and protease inhibitor cocktail (Sigma). The cell lysates were sonicated for 10-15 sec on ice, followed by centrifugation at 14,000 rpm for 15 min at 4°C and the supernatant was then collected. For tissue protein isolation, the liver tissue samples were washed with PBS and homogenized in 1.0 ml RIPA buffer using a micropestle (Tarsons) in a 1.5 ml microcentrifuge tube. The homogenized samples were sonicated, centrifuged at 14,000 rpm and the supernatant was stored in aliquots at −80°C until further use. The protein concentrations in the cell lysates or tissues were estimated by Bradford Assay (Bio-Rad). Equal amount of protein samples (~30-50 μg) per condition were separated by sodium dodecyl sulphate polyacrylamide gel electrophoresis (SDS-PAGE) gel and transferred to activated PVDF membrane (Pall Life Sciences). After blocking with 2-5% of BSA/ non-fat milk for 1 hour at room temperature, the membranes were incubated with specific primary antibody [HMGCR (Abcam, ab174830) at 1:1000 dilution, β-Actin (Sigma, A5441) at 1:7500 dilution, Vinculin (Sigma, V9131) at 1:7500 dilution, Egr1 (CST, 4153) at 1:1000 dilution] overnight at 4°C. After washing with 1xTBST, the membrane was incubated with HRP-conjugated secondary antibody specific for either rabbit (BioRad # 170-6515 at 1:1500 dilution for HMGCR or Egr1) or mouse (Jackson Immunoresearch # 115-035-003 at 1:5000 dilution for β-actin) for 1 hr. The protein bands were detected using Clarity™ Western ECL Substrate kit (Bio-Rad) and signal was captured by Chemidoc XRS+ Chemiluminescence Detection system (Bio-Rad). Densitometric analysis of Western blots was performed using Image Lab (Bio-Rad) or NIH ImageJ software [23]. All the Western blot experiments were repeated at least three times and representative images are shown.

### Chromatin Immunoprecipitation (ChIP) Assays

AML12 cells, at 60-80% confluency, were cross-linked using formaldehyde at room temperature for 12 min. Next, chromatin was isolated and sheared by sonication followed by a pre-clearing step with Rec protein G-Sepharose 4B beads (Invitrogen). Immunoprecipitation reactions of the pre-cleared samples were carried out by incubation with 5 μg each of ChIP grade antibodies, i.e. anti-Egr1 and pre-immune anti-rabbit IgG (Sigma, I5006) overnight at 4°C. The immunoprecipitated samples were captured by Rec protein G-Sepharose 4B beads, eluted, reverse cross-linked and purified by phenol-chloroform extraction. qPCR was carried out using equal amount of the purified chromatin as template to amplify two different DNA regions harboring Egr1 binding sites in the proximal (~500 bp) promoter domain of miR-27a promoter using two primer pairs (Fig.5A) (P1-FP: 5’-TCAAGATAGGCAGGCAAGC-3’ and P1-RP: 5’-AGCACAGGGTCAGTTGGAAA-3’; P2-FP: 5’-TTTGTAGGGCTGGGGTAGAG-3’ and P2-RP: 5’-CTGATCCACACCCTAGCCC-3’). Results were expressed as fold enrichment over IgG signal or background.

### Data presentation and Statistical analysis

All transient transfection experiments were performed at least three times and results were expressed as mean ± SEM of triplicates. Prism 5 program (GraphPad Software, USA) was used to determine the level of statistical significance by Student’s t-test or one-way ANOVA with Newman-Keuls’s post-test, as appropriate.

## RESULTS

### Comparative genomics analysis of mouse and rat *Hmgcr* gene sequences

Analysis of rat elevated lipid/cholesterol-QTLs on *Hmgcr*-harboring chromosome 2 within the range of 26000000-28000000 bp detected six lipid-related QTLs (Fig.S1A); these QTLs and respective LOD [logarithm (base 10) of odds] scores were retrieved from the Rat Genome Database and are shown in Table S1. Interestingly, among the QTLs harboring the *Hmgcr* locus (27480226-27500654 bp, RGD ID: 2803), Stl27 (23837491-149614623 bp) and Stl32 (22612952-67612952 bp) displayed high linkage (LOD score=4.4, 3.2, respectively) with blood triglyceride levels (Fig.S1A) while Scl55 (26186097-142053534 bp) showed significant linkage with blood cholesterol levels (LOD score=2.83). Moreover, alignment of mouse and rat *Hmgcr* locus using mVISTA showed >75% homology between these rodents at exons, introns and untranslated regions (Fig.S1B). Interestingly, the extent of homology between the twenty *Hmgcr* exons in mouse and rat was higher (>85%) than the noncoding regions (Fig.S1B). Thus, mouse *Hmgcr* appeared as a logical candidate gene for studying the mechanisms of hypercholesterolemia.

### Identification of potential miRNAs involved in *Hmgcr* regulation

Since miRNA prediction tools employ different algorithms, their outputs may vary and often a set of miRNAs predicted by one tool may not overlap with the others. Hence, in this study we used multiple tools to predict the miRNA binding sites in the 3’-UTR of *Hmgcr* to increase the accuracy of target predictions. *In silico* predictions followed by extensive screening procedures shortlisted 7 miRNAs (miR-27a, miR-27b, miR-28, miR-124, miR-345, miR-351 and miR-708; Table S4). Since the miR-27a, miR-27b, miR-28 and miR-708 binding sites are highly conserved across mammals including humans, we examined their expression profile in different human tissues using DASHR database. The expression of HMGCR transcript levels in different human tissues was mined from GTEx portal. Interestingly, *HMGCR* expression showed a significant inverse correlation with the miR-27a expression (Fig.1D) (Pearson *r* = −0.9007, p<0.05). In contrast, miR-27b, miR-28 and miR-708 levels did not exhibit an inverse correlation with *HMGCR* expression in these tissues (Fig.S2). Since the expression of miR-124, miR-345 and miR-351 across various tissues was near/below the detectable range for functional repression of target genes (viz., below 100 RPM) [10], these were not considered for correlation analysis. Moreover, only miR-27a and miR-27b have been validated to interact with Hmgcr via HITS-CLIP analysis as reported in the TarBase database (Table S2). All these lines of evidence indicate a possible role of miR-27a and miR-27b in the post-transcriptional regulation of Hmgcr. Hence, miR-27a and miR-27b were selected to further investigate their interactions with Hmgcr.

**Figure 1.**
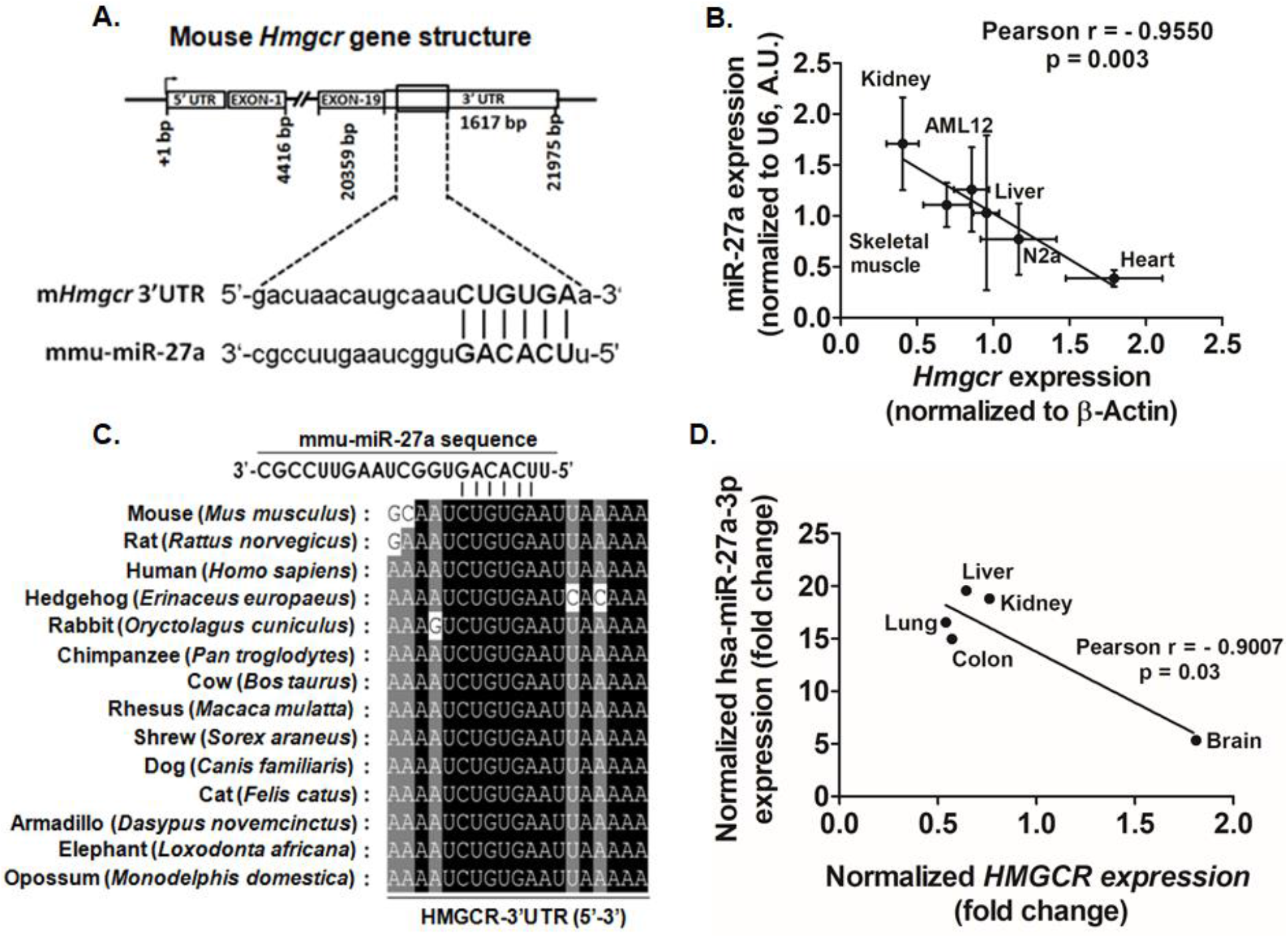
miR-27a binding sites in the 3’-UTR of *Hmgcr* and inverse correlation between mmu-miR-27a-3p/hsa-miR-27a-3p and *Hmgcr/HMGCR* expression. (A) Schematic representation of *Hmgcr* showing the miR-27a binding site in the 3’-UTR. The complementarity between the seed region of miR-27a and mHmgcr-3’UTR is depicted in bold and capital. (B) Negative correlation between *Hmgcr* mRNA and miR-27a expression in cultured AML12, N2a cells, rat tissues (number of biological/technical replicates was at least three in each case). (C) The conservation of the miR-27a binding site in the 3’-UTR of *Hmgcr* across different mammals. (D) Inverse correlation between *HMGCR* and hsa-miR-27a-3p expression in different human tissues.

### Direct interaction of miR-27a with *Hmgcr* 3’-UTR down-regulates Hmgcr protein level in hepatocytes

To examine whether miR-27a directly interacts with *Hmgcr*, miR-27a expression plasmid was co-transfected with m*Hmgcr* 3’UTR/luciferase construct into AML12 and HuH-7 cells, respectively (Fig.2A, B). Indeed, miR-27a over-expression caused a significant dose-dependent reduction in the m*Hmgcr* 3’UTR reporter activity in both AML12 (up to ~85%, p<0.001) and HuH-7 cells (up to ~34%, p<0.01). In contrast, co-transfection of m*Hmgcr* 27a mut 3’UTR construct (devoid of miR-27a binding site) with miR-27a expression plasmid showed no significant change in 3’UTR reporter activity (Fig.2A, B). Further, qPCR analysis showed that over-expression of pre-miR-27a led to a dose-dependent increase in miR-27a expression in AML12 (up to ~1770%, p<0.01) and HuH-7 (up to ~125%, p<0.01) cells. Consistent with the m*Hmgcr* 3’UTR reporter activity, over-expression of miR-27a caused a decrease in endogenous Hmgcr/HMGCR protein level (Fig.2C, D).

**Figure 2.**
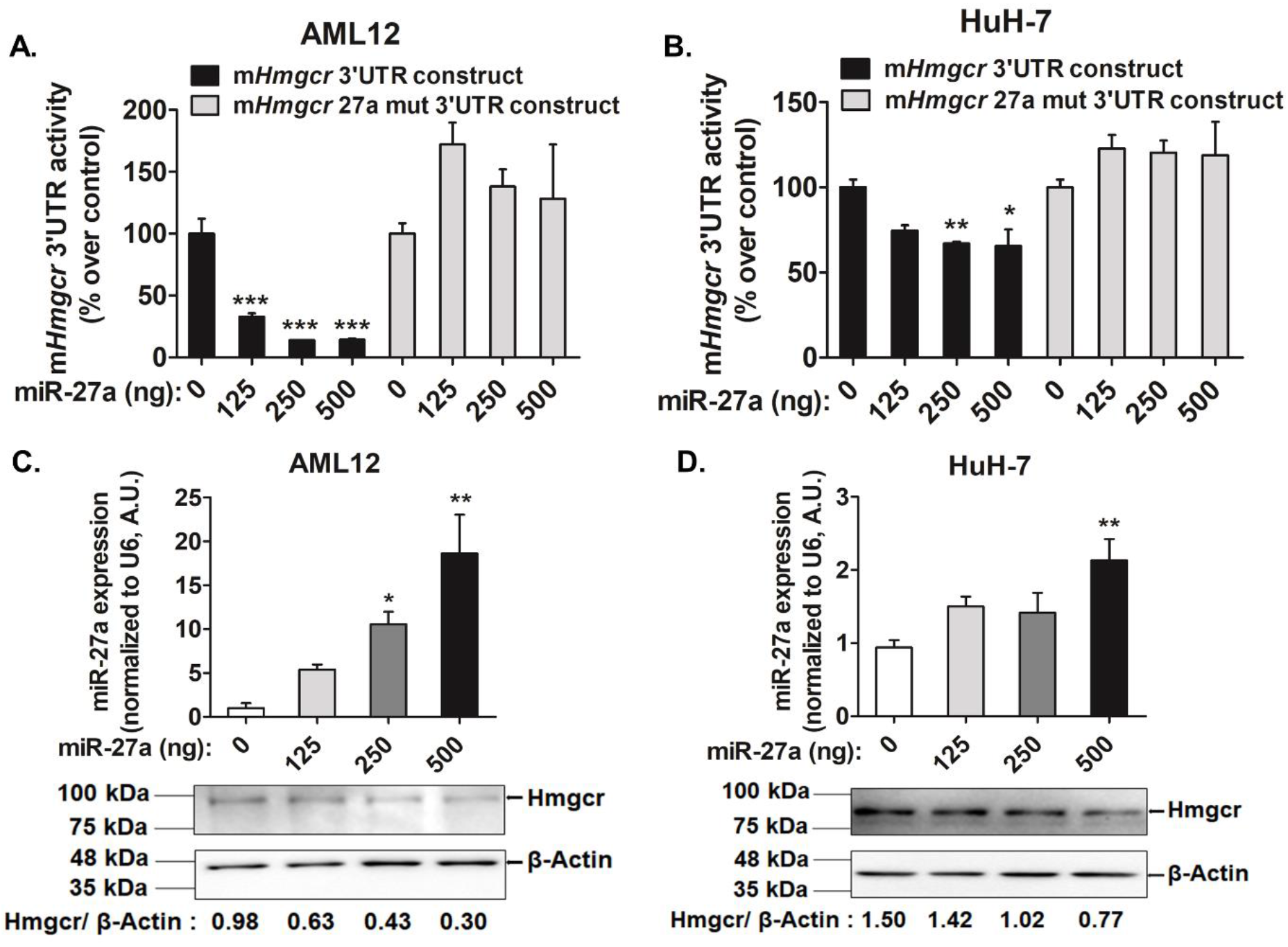
miR-27a negatively regulates Hmgcr expression in cultured hepatocytes. The m*Hmgcr* 3’UTR reporter construct or m*Hmgcr* 27a mut 3’UTR construct (500 ng) was co-transfected with miR-27a expression plasmid in (A) AML12, (B) HuH-7 cells followed by luciferase assays (n=5). The results are mean ± SEM of triplicate values. Transfection of miR-27a expression plasmid results in over-expression of miR-27a in (C) AML12 and (D) HuH-7 cells, respectively (n=5). Statistical significance was determined by one-way ANOVA with Newman-Keuls multiple comparison test. *p<0.05, **p<0.01, ***p<0.001 as compared to control. Hmgcr down-regulation was confirmed by Western blot analysis (C and D) (n=3).

Despite the predicted binding site of miR-27b in the *Hmgcr* 3’-UTR, co-transfection of miR-27b expression plasmid did not cause significant decrease in m*Hmgcr* 3’UTR reporter activity (Fig.S3A). Moreover, to test the specificity of interactions of miR-27a with *Hmgcr* 3’-UTR, we carried out co-transfection experiments with miR-764 which does not have binding sites in the *Hmgcr* 3’-UTR region; no change in the *Hmgcr* 3’-UTR activity was observed (Fig.S3B).

Furthermore, transfection of locked nucleic acid inhibitor of miR-27a (LNA 27a) in AML12 cells diminished the endogenous miR-27a levels (by ~15-fold, p<0.05; Fig.3A) and enhanced Hmgcr mRNA (by ~3.8-fold, p<0.05, Fig.S4A)/protein levels (Fig.3B). On the other hand, transfection of miR-27a mimic showed enhanced miR-27a levels (by ~164-fold, p<0.05; Fig.3C) and diminished Hmgcr mRNA (Fig.S4B)/protein levels (Fig.3D). In corroboration, RIP (Ribonucleoprotein immunoprecipitation) assays with antibody specific to Ago2 (an integral component of RNA induced-Silencing Complex, RISC) showed enrichment of *HMGCR* transcript level (by ~3.5-fold, p<0.05) in the Ago2-immunoprecipitated RNA fraction of HuH-7 cells over-expressing miR-27a as compared to the control condition, thereby confirming the *in vivo* interaction of miR-27a with *Hmgcr* 3’-UTR in the context of RISC (Fig.3E). RNA fraction immunoprecipitated using pre-immune anti-mouse IgG antibody (control) showed no significant difference in *HMGCR* levels between miR-27a over-expression and basal conditions (Fig.3E).

**Figure 3.**
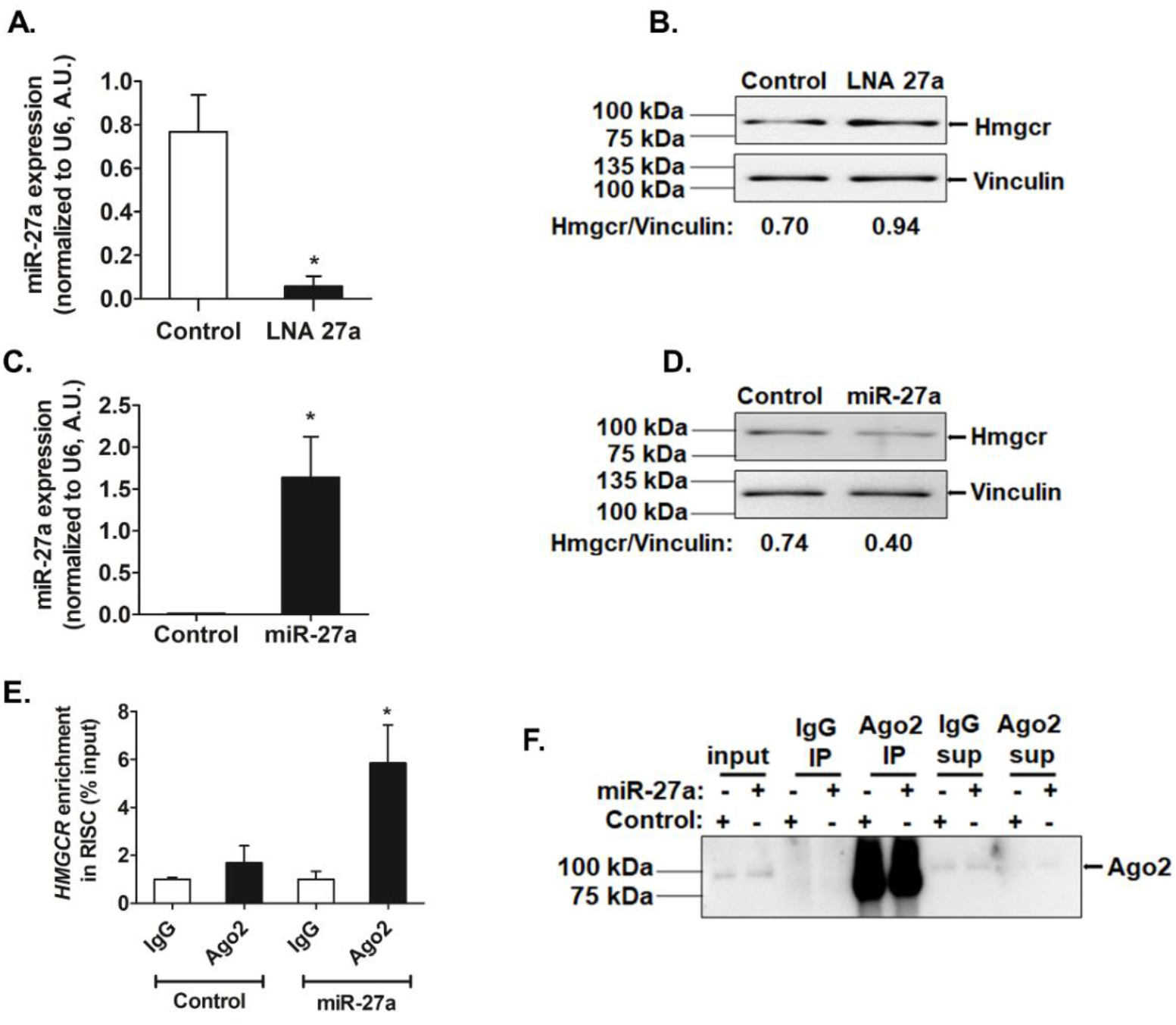
Specific interactions of miR-27a with *Hmgcr* 3’-UTR. The relative expression of (A) miR-27a upon transfection of 60 nM of control oligo or locked nucleic acid inhibitor of miR-27a (LNA27a) in AML12 cells was determined by qPCR (n=3). (B) Representative Western blot analysis of Hmgcr protein levels in AML12 cells upon transfection of 60 nM of control oligo or LNA27a (n=3). (C) miR-27a levels upon transfection of 1 μg of control oligo or miR-27a mimic in AML12 cells were determined by qPCR (n=3). (D) Representative Western blot analysis of Hmgcr protein levels in AML12 cells upon transfection of 1 μg of control oligo or miR-27a mimic (n=3). Statistical significance was determined by Student’s *t*-test (unpaired, 2-tailed). *p<0.05 as compared to control. (E) Ago2 Ribonucleoprotein precipitation analysis in HuH-7 cells over-expressing miR-27a. The *HMGCR* enrichment was normalized to corresponding input in each condition; expressed as % input and is mean ± SEM for triplicates (n=3). (F) Ago2 immunoprecipitation was confirmed by Western blotting (n=3). Statistical significance was determined by one-way ANOVA with Newman-Keuls multiple comparison test. *p<0.05 with respect to control Ago2 condition.

### Role of intracellular cholesterol level in miR-27a-mediated regulation of Hmgcr

Because Hmgcr protein level is dependent on intracellular cholesterol level and regulated by a negative feed-back mechanism, we sought to determine whether modulating endogenous cholesterol level affects the post-transcriptional regulation of *Hmgcr*. Accordingly, AML12 cells were treated with different concentrations of either cholesterol (5, 10 and 20 μg/ml) or methyl-β-cyclodextrin (MCD, an oligosaccharide that reduces intracellular cholesterol level; 1, 2.5 and 5 mM) followed by Western blot analysis for Hmgcr (Fig.4A, B). Since 20 μg/ml of cholesterol or 5 mM of MCD showed effective reduction or augmentation in Hmgcr protein level, respectively, these doses were used for further experiments. Interestingly, cholesterol treatment showed ~1.9-fold (p<0.05) enhancement of miR-27a level (Fig.4C) while cholesterol depletion caused a ~3-fold (p<0.05) reduction in endogenous miR-27a level (Fig.4D). Increase/decrease in the intracellular cholesterol level was confirmed by Filipin staining (Fig.4E, F). Further, Ago2-RIP assays in cholesterol-treated HuH-7 cells revealed significant enrichment of *HMGCR* (~2.2-fold, p<0.01; Fig.4G) and miR-27a (~1.7-fold, p<0.001; Fig.4H) suggesting *in vivo* interaction of *HMGCR* with miR-27a under elevated cholesterol conditions.

**Figure 4.**
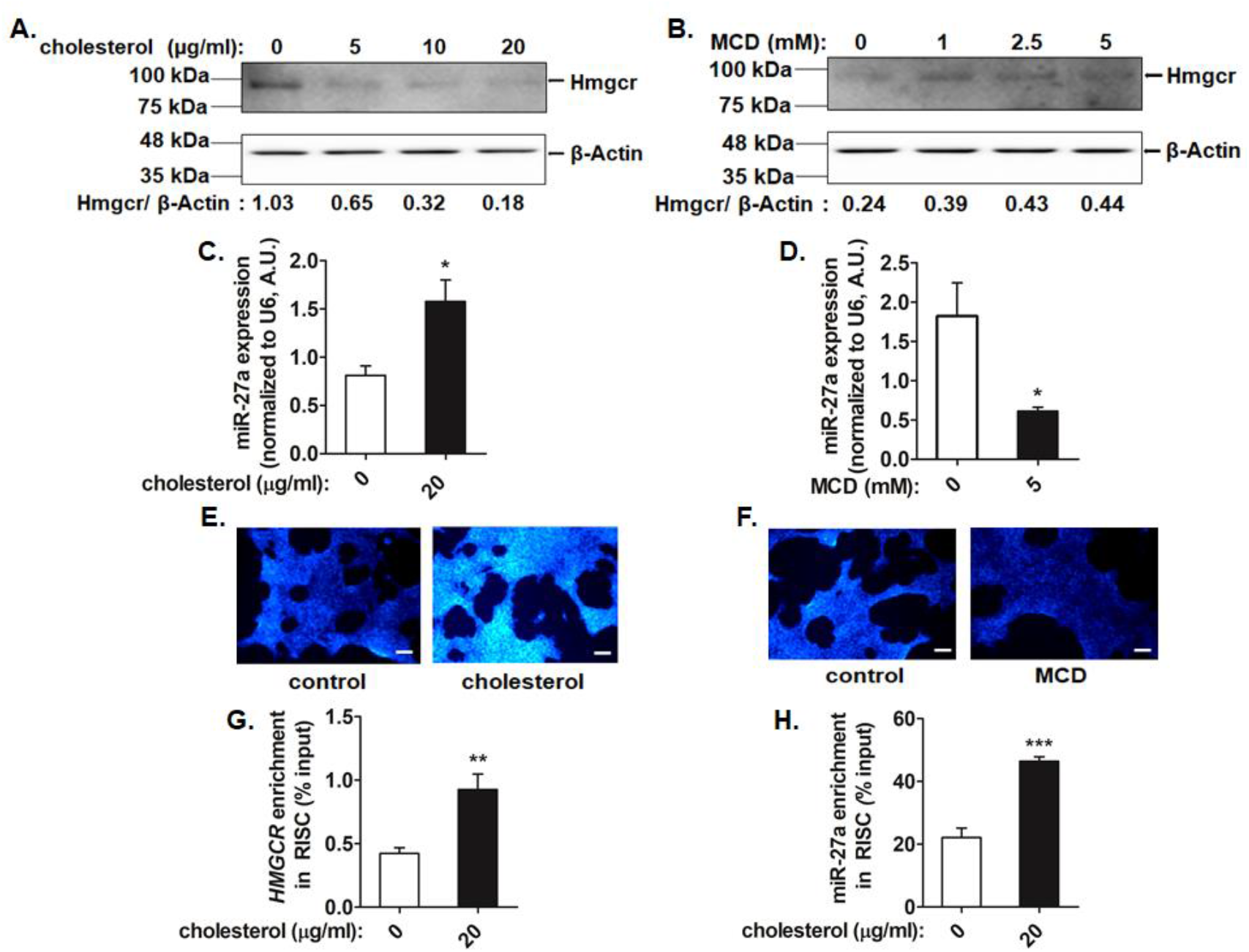
Intracellular cholesterol modulates the expression of miR-27a and augments the interactions of miR-27a with the *Hmgcr* 3’-UTR. AML12 cells were treated with increasing doses of either cholesterol or cholesterol-depleting reagent methyl-β-cyclodextrin (MCD) for 6 hrs or 15 minutes, respectively. After incubation for 6-9 hrs in serum free media, the cells were processed for RNA, protein isolation and fluorescence microscopy. Western blot analysis of total protein isolated from AML12 cells treated with (A) cholesterol or (B) MCD was carried out for Hmgcr and β-actin levels (n=3). miR-27a levels were also determined by qPCR on (C) exogenous cholesterol (20 μg/ml) or (D) MCD (5 mM) treatment in AML12 cells (n=3). The miR-27a expression was normalized to U6 RNA and indicated as mean ± SEM for triplicate values. (E) Cholesterol or (F) MCD treatments in AML12 cells were confirmed by staining intracellular cholesterol using Filipin stain followed by fluorescence microscopy. Scale bar: 170 μm. (G and Enrichment of *HMGCR* and miR-27a upon exogenous cholesterol treatment in Ago2-immunoprecipitated RNA from HuH-7 cells. HuH-7 cells were treated with 20 μg/ml of cholesterol for 6 hrs and the total RNA fraction from the Ago2/IgG-immunoprecipitated samples (in basal and cholesterol-treated cells) was subjected to qPCR using (G) *HMGCR* and (H) miR-27a specific primers (Table S3; n=3). The *HMGCR*/miR-27a enrichment was normalized to the corresponding input in each condition and represented as % input. Statistical significance was determined by Student’s *t*-test (unpaired, 2-tailed). *p<0.05, **p<0.01 and ***p<0.001 as compared to the basal condition.

### Role of Egr1 in miR-27a expression under basal and modulated intracellular cholesterol conditions

To understand the possible mechanism of miR-27a regulation, we predicted transcription factor binding sites in the miR-27a promoter domain (~500 bp) using two programs: LASAGNA and JASPAR (Table S2). Egr1, a transcription factor (predominantly expressed in the liver) that plays a crucial role in the transcriptional regulation of most cholesterol biosynthesis genes including *Hmgcr* [24], had six putative binding sites as two separate clusters in the miR-27a promoter domain (Fig.5A).

**Figure 5.**
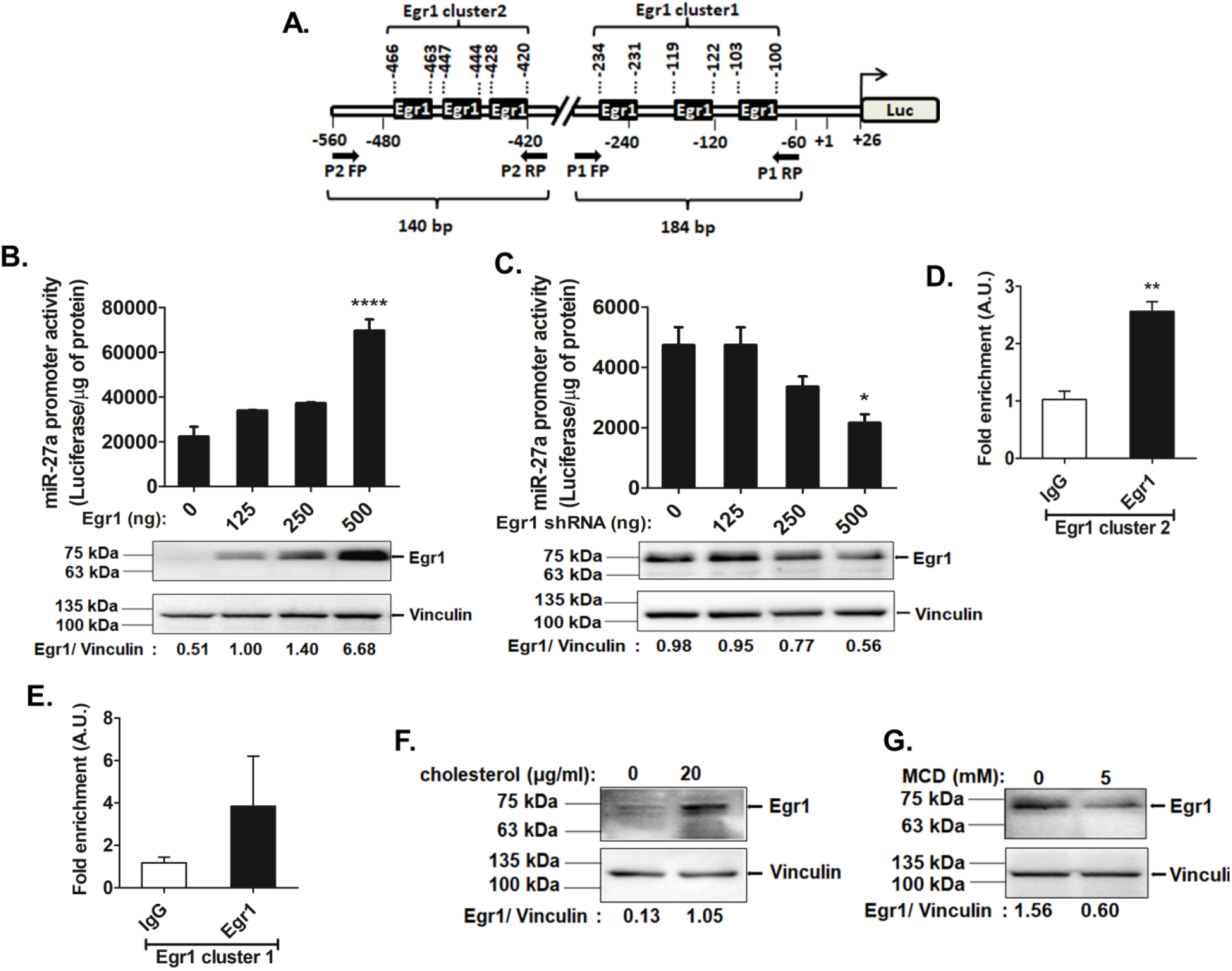
Egr1 modulates miR-27a expression under basal and altered cholesterol conditions. (A) Schematic representation of the proximal mmu-miR-27a promoter domain harboring multiple Egr1 binding sites. AML 12 cells were co-transfected with either (B) Egr1 expression plasmid or (C) Egr1 shRNA expression plasmid and miR-27a promoter construct (500 ng) (n=5). Results are expressed as mean ± SEM for triplicate values. Statistical significance was determined by one-way ANOVA with Newman-Keuls multiple comparison test. *p<0.05, ****p <0.0001 as compared to basal condition. Over-expression/down-regulation of Egr1 was confirmed by Western blot analysis (n=3). ChIP assays using Egr1/ IgG immunoprecipitated chromatin isolated from AML12 cells. qPCR was performed with immunoprecipitated DNA using primer pair (D) P2 (amplifying Egr1 cluster 2) and (E) P1 (amplifying Egr1 cluster in miR-27a promoter (n=3). Statistical significance was determined by Student’s *t*-test (unpaired, 2-tailed). *p<0.05 and **p<0.01 as compared to control. AML12 cells were treated with either (F) 20 μg/ml of cholesterol [(3β)-cholest-5-en-3-ol] or (G) 5 mM of cholesterol-depleting reagent methyl-β-cyclodextrin (MCD) for 6 hrs or 15 minutes, respectively; Western blots quantifying Egr1 levels are shown (n=3).

Next, we validated the role of Egr1 in miR-27a expression by co-transfection experiments. Egr1 over-expression caused a ~3-fold enhancement of miR-27a promoter activity (p<0.0001) (Fig.5B) while Egr1 down-regulation resulted in a ~2.2-fold reduction in miR-27a promoter activity (p<0.05; Fig.5C). A concomitant increase (~4-fold, p<0.05)/decrease (~2-fold, p<0.05) in the endogenous miR-27a levels upon Egr1 over-expression/down-regulation was also observed (Fig.S5). Thus, Egr1 may play a crucial role in the transcriptional activation of miR-27a. Indeed, ChIP (Chromatin Immunoprecipitation) assays confirmed the *in vivo* interaction of Egr1 with the miR-27a proximal (~-500 bp) promoter domain. qPCR analysis using Egr1 antibody-immunoprecipitated chromatin revealed ~2.5-fold enrichment of the miR-27a promoter domain with P2 primer pair (p<0.001; Fig.5D) whereas no significant fold-enrichment of the miR-27a promoter domain was observed using primer pair P1 (Fig.5E) suggesting that the Egr1 sites predicted in this domain (Egr1 cluster 1) may not be functional under basal conditions. Interestingly, Western blot analysis revealed that exogenous cholesterol treatment augmented Egr1 level (Fig.5F) while its level diminished upon cholesterol depletion (Fig.5G) suggesting that intracellular cholesterol level may regulate miR-27a expression via Egr1. Furthermore, downregulation of Egr1 abrogated the cholesterol-mediated activation of miR-27a promoter activity (Fig.S6) suggesting that Egr1 plays a key role in regulating miR-27a expression under modulated cholesterol levels.

### miR-27a mimic reduces hepatic Hmgcr expression and plasma cholesterol levels in high cholesterol diet-fed *Apoe*^−/−^ mice

To test if miR-27a mimics can modulate plasma lipid levels in a model of hyperlipidemia, *Apoe*^−/−^ mice on a high cholesterol diet regimen for 10 weeks were injected with either saline or 5 mg/kg dose of miR-27a mimic or control oligo as lipid emulsion formulations (Fig.6A). Tissue analysis of miR-27a levels revealed ~4-fold (p<0.05), ~5.8-fold (p<0.05) and ~4.3-fold (p<0.05) up-regulation of miR-27a expression in the liver, heart and adipose tissues, respectively, in miR-27a mimic-injected animals as compared to the control oligo group. However, miR-27a levels did not differ in the kidney and brain tissues of miR-27a mimic group as compared to control group (Fig.6B) suggesting efficient delivery of the miR-27a mimic to the liver, heart and adipose tissues.

**Figure 6.**
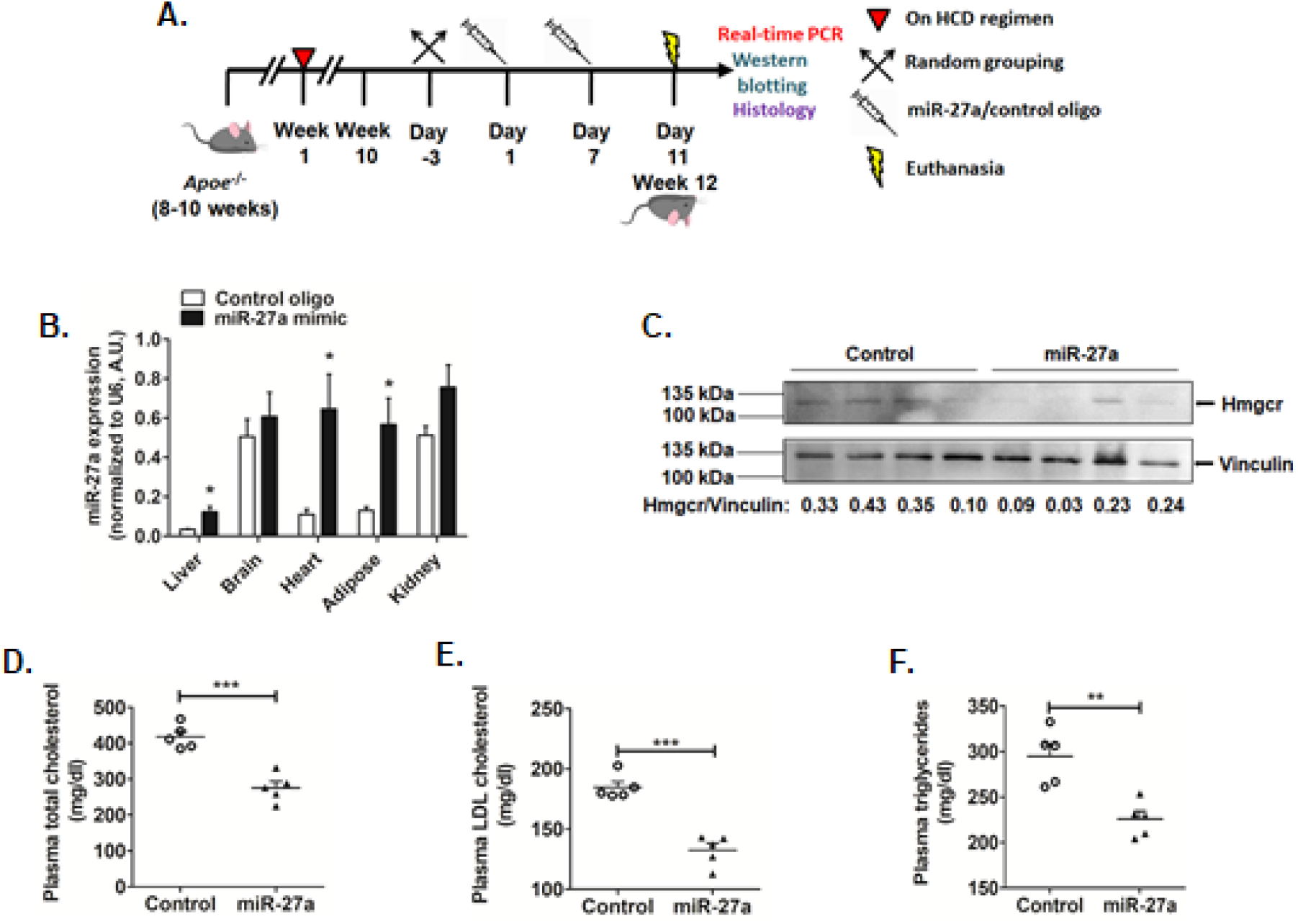
miR-27a diminishes Hmgcr and plasma lipid levels in high cholesterol diet-fed *Apoe*^−/−^ mice. (A) Male *Apoe*^−/−^ mice (8-10 weeks old) were fed a high-cholesterol diet (HCD) regimen for 10 weeks and injected with either miR-27a mimic/control oligo twice over a period of 11 days and then euthanized. (B) qPCR for miR-27a levels in various tissues. (C) Western blot analysis of Hmgcr protein in *Apoe*^−/−^ mice liver tissues representative of three experiments. Plasma levels of (D) total cholesterol, (E) LDL cholesterol and (F) triglycerides from overnight-fasted animals. Mean ± SEM, n=5-6 animals per group. Statistical significance was determined by Student’s *t*-test (unpaired, 2-tailed). *p<0.05, **p<0.01 and ***p<0.001 as compared to control group.

Western blot analysis in liver tissues of these mice revealed diminished Hmgcr protein level in miR-27a mimic-injected mice as compared to control (Fig.6C, S7A). The anti-Hmgcr antibody detected a ~120 kDa band suggesting predominantly glycosylated form of the protein in these tissues. The hepatic cholesterol level in miR-27a-injected animals showed a modest decrease, which did not differ significantly in comparison to the control group (Fig.S7C).

Next, we assessed the effect of miR-27a mimics on the circulating lipid levels. These parameters were measured 4 days after the second injection of miR-27a mimic/control oligo. Interestingly, the miR-27a mimic group showed reduced plasma total cholesterol (~1.5-fold, p<0.001), LDL cholesterol levels (~1.4-fold, p<0.001) and triglycerides (~1.3-fold, p<0.01) (Fig.6D, E, F) as compared to the control group. In contrast, miR-27a mimic enhanced the plasma HDL cholesterol levels (~1.5-fold, p<0.05) (Fig.S7B) in comparison to the control group.

It is important to note that plasma ALT, AST and CK levels were not elevated in this model indicating no obvious liver/muscle injury upon miR-27a mimic injections (Fig.S7D, E, F). These findings suggest the therapeutic potential of miR-27a mimic as a safe and efficient cholesterol lowering agent.

### miR-27a represses Hmgcr by post-translational inhibition followed by mRNA degradation

In order to unfold the mechanism of action of miR-27a on *Hmgcr*, mRNA stability assays using actinomycin D were performed in AML12 cells over-expressing miR-27a (Fig.S8). Interestingly, no significant change in Hmgcr mRNA half-life was observed upon miR-27a over-expression till 24 hours (Fig.S8A, B). In contrast, the Hmgcr protein levels showed a time-dependent reduction following actinomycin D treatment in AML12 cells transfected with miR-27a (Fig.S8C, D). In corroboration, over-expression of miR-27a did not alter steady state Hmgcr mRNA level (Fig.S8E). However, 36 hours post-transfection of LNA 27a in AML12 cells, endogenous Hmgcr mRNA levels were significantly enhanced (Fig.S4A). These observations suggest that Hmgcr repression by miR-27a is mediated predominantly by translational attenuation followed by mRNA degradation.

### miR-27a targets multiple genes in the cholesterol regulatory pathways

In line with our observations and reported role of miR-27a in lipid metabolism, we tested if miR-27a targets other genes crucial for cholesterol homeostasis. Interestingly, *in silico* predictions and PANTHER pathway analysis revealed six additional cholesterol biosynthesis-related gene targets: 3-Hydroxy-3-Methylglutaryl-CoA synthase 1 (*Hmgcs1*), Mevalonate kinase (*Mvk*), Diphosphomevalonate decarboxylase (*Mvd*), Geranylgeranyl pyrophosphate synthase (*Ggps1*), Squalene synthase (*Fdft1*), Squalene epoxidase (*Sqle*) (Fig.7A). Indeed, miR-27a mimic treatment in AML12 cells or high cholesterol diet-fed *Apoe*^−/−^ mice diminished the expression of *Mvk*, *Fdft1*, *Sqle*, *Ggps1* and *Mvd* (Figs.9B, C). In addition, miR-27a augmentation also resulted in diminished expression of lipoprotein uptake-related genes viz. low density lipoprotein receptor (*Ldlr*) and Scavenger Receptor Class B Member1 (*Scarb1*) (Fig.S4C, D). Thus, miR-27a seems to play a key role in the global regulation of genes involved in cholesterol homeostasis.

**Figure 7.**
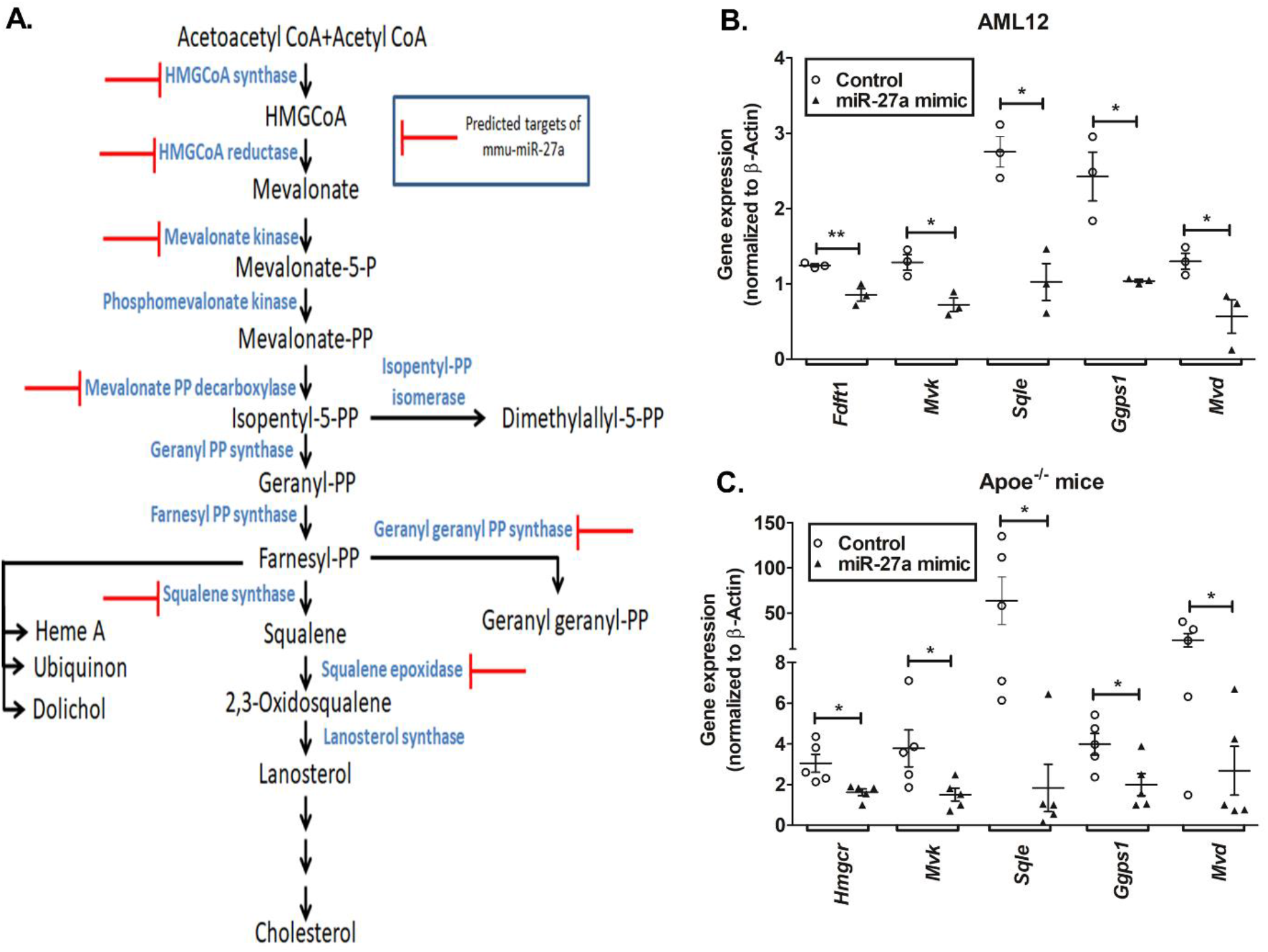
Augmentation of miR-27a represses multiple genes in the cholesterol biosynthesis pathway. (A) Potential targets of miR-27a in the cholesterol biosynthesis pathway predicted by miRWalk, RNAhybrid and TargetScan were categorized based on their molecular functions using PANTHER database. Target genes are indicated by red bar-headed lines. qPCR analysis of predicted genes in (B) AML12 (n=3), (C) liver tissues of high cholesterol diet-fed *Apoe*^−/−^ mice transfected/injected with miR-27a mimic/control oligo (n=5-6 animals per group). Statistical significance was determined by Student’s *t*-test (unpaired, 2-tailed). *p<0.05, **p<0.01 and ***p<0.001 as compared to control group.

## DISCUSSION

### Overview

In view of the important role of Hmgcr in cholesterol biosynthesis its potential regulation by microRNAs has been of major interest in recent years. Notably, miR-29a, miR-185, miR-150, miR-548p, miR-21, miR-195 and miR-342 have so far been reported to interact with the Hmgcr transcript [25–31]. We undertook extensive computational and experimental analyses to identify the key miRNAs that may regulate *Hmgcr* gene expression under basal as well as pathophysiological conditions. Our *in vitro* data provides several lines of evidence for regulation of *Hmgcr* expression by miR-27a (Figs. 1, 2 and 3). We also observed the effects of miR-27a mimic to down-regulate hepatic expression of Hmgcr and plasma lipid levels in a high cholesterol diet-fed atherosclerotic mouse model (Figs.6, S7).

### Pathophysiological implications of Hmgcr regulation by miR-27a

HMGCR is tightly regulated by sterols via transcriptional, post-transcriptional and post-translational mechanisms [32]. In brief, elevated sterols diminish *HMGCR* expression by inhibiting Sterol Regulatory Element Binding Protein 2 (SREBP-2) transcription factor [33]. The post-transcriptional and post-translational regulatory systems operate independent of SREBP pathway and form an important aspect of sterol-mediated HMGCR regulation. Post-translational regulation is executed by sterol or non-sterol intermediates via INSIG dependent ER-associated protein degradation (ERAD) mechanism involving ubiquitin-proteasomal degradation of HMGCR [7]. However, the effect of elevated sterols on miRNA-mediated *HMGCR* regulation is partially understood. In view of crucial role of sterols in HMGCR regulation, do enhanced cholesterol levels modulate endogenous miR-27a levels? Indeed, cholesterol treatment to AML12 cells enhanced miR-27a level and diminished Hmgcr protein level (Fig.4A, C). In order to rule out the possibility that this repression in Hmgcr protein level is solely because of INSIG-mediated ERAD mechanism, we performed RIP assays in HuH-7 cells treated with cholesterol. Our Ago2-RIP assays further confirmed enhanced interaction of miR-27a with *HMGCR* under elevated cholesterol level suggesting that post-transcriptional regulation of Hmgcr by miR-27a is an additional mechanism for Hmgcr repression under high cholesterol conditions (Fig.4G, H). Thus, this study revealed the crucial role of intracellular cholesterol on Hmgcr expression via miR-27a. This is in corroboration with enhanced miR-27a level in liver tissues of *Apoe*^−/−^ mice fed with a high cholesterol diet regimen as compared to *Apoe*^−/−^ mice fed with a normal chow diet (Fig.S9).

Are there other microRNAs that modulate lipid or lipoprotein synthesis/secretion into the circulation? There are limited reports on the regulatory roles of miRNAs in cholesterol homeostasis. miR-122 inhibition in mice results in diminished plasma cholesterol and triglyceride levels with reduced fatty acid and cholesterol synthesis [34]. Interestingly, deletion of miR-122 in mice causes enhanced hepatic lipid, plasma alanine aminotransferase and alkaline phosphatase levels with the development of steatohepatitis, fibrosis and hepatocellular carcinoma (HCC) with age [35]. Likewise, miR-34 over-expression in mice reduced plasma triglyceride and cholesterol levels but augmented hepatic triglyceride levels causing hepatosteatosis [36]. Recent studies indicate that miR-30c lowers plasma cholesterol in different mice models of atherosclerosis, hypercholesterolemia and metabolic disorders without increasing plasma transaminases or causing hepatic steatosis [19,37]. These studies highlight the potential of miRNAs as therapeutic agents in the treatment of hypercholesterolemia.

For validation of our *in vitro* findings, we tested the effect of miR-27a mimic in high cholesterol diet-fed *Apoe*^−/−^ mice. An improved lipid-based formulation was employed for the efficient delivery of the miR-27a mimic via tail vein injection to major organs viz. the liver, heart and adipose tissues as confirmed by the tissue analysis of miR-27a levels (Fig.6B). Administration of miR-27a over a period of 11 days induced a significant reduction in the plasma total cholesterol, LDL cholesterol and triglyceride levels with a concomitant increase in the HDL cholesterol levels (Fig.6D, E, F, Fig.S7B).

miR-27a appears to diminish the plasma cholesterol levels by reducing the Hmgcr protein level in the liver (Fig.6C, S7A). The reductions in the plasma triglycerides are consistent with a previous report wherein miR-27a expression diminished triglyceride levels in mouse hepatocytes mainly by targeting fatty acid synthase (Fas) and stearyl CoA desaturase 1 (Scd1) [38]. We speculate that the increase in plasma HDL cholesterol level might result from the miR-27a-mediated regulation of Scarb1, an HDL cholesterol receptor (Fig.S4D). It is interesting to note that the food-derived remnant cholesterol has a modest contribution to the total cholesterol level. Moreover, it has been previously reported that remnant lipoproteinemia in APOE ^−/−^ minipigs is not efficient in initiating atherosclerosis but contributes to the atherosclerotic progression of pre-existing lesions [39]. Therefore, miR-27a mimic may diminish the overall plasma total cholesterol levels in high cholesterol diet-fed Apoe^−/−^ mice by modulating the LDL, HDL and other cholesterol fractions.

Does the miR-27a mimic treatment cause any adverse effects in the diet-induced atherosclerosis mouse model? We did not detect any significant changes in plasma ALT, AST and CK levels suggesting that the animals did not suffer from any liver or muscle injury (Fig.S7D, E, F). However, additional studies are required to test the long-term effects of the miR-27a mimic to further evaluate its therapeutic potential

### Molecular mechanisms of Hmgcr regulation by miR-27a

Generally, miRNAs exert their action by translational inhibition followed by mRNA deadenylation, decapping and decay [40]. Our actinomycin D chase experiments showed that Hmgcr mRNA half-life did not change whereas the Hmgcr protein level diminished in a time-dependent manner (Fig.S8). Consistently, the steady-state Hmgcr mRNA level in hepatocytes did not diminish upon miR-27a over-expression (Fig.S8). However, Hmgcr mRNA levels were significantly altered at time points exceeding 24 hours of transfection in hepatocytes or miR-27a mimic injection in *Apoe*^−/−^ mice (Fig.S4A, Fig.7C). These observations are consistent with a previous report which suggests that miRNAs initially reduce target protein levels without affecting mRNA levels but diminish mRNA levels at later time points [41]. Therefore, we speculate that miR-27a-mediated Hmgcr repression mainly involves translational control followed by mRNA degradation.

In view of the key role of miR-27a in regulating Hmgcr, we sought to unravel how miR-27a might be regulated under basal and pathophysiological conditions. Computational and experimental analyses suggested a crucial role of Egr1 in the activation of miR-27a expression. Egr1, a zinc finger transcription factor belonging to the early growth response gene family, binds to a GC-rich consensus region [42] and regulates genes involved in physiological stress response, cell metabolism, proliferation and inflammation [43]. Egr1, majorly expressed in the liver, targets multiple cholesterol biosynthesis genes [24,44]. In light of putative Egr1 binding sites in the proximal miR-27a promoter domain (Fig.5A), we investigated the role of Egr1 in miR-27a expression. Indeed, over-expression/down-regulation of Egr1 resulted in enhanced/diminished miR-27a promoter activity in AML12 cells (Fig.5B, C). ChIP assays also confirmed the *in vivo* interaction of Egr1 with the miR-27a promoter (Fig.5D). Therefore, our results suggest that Egr1 regulates miR-27a expression under basal conditions.

### Role of miR-27a in global regulation of cholesterol homeostasis

miR-27a, an intergenic miRNA, is transcribed from the miR-23a-miR-27a-miR-24 cluster located on chromosome 8 in mice. Dysregulation of miR-27a has been associated with several cardiovascular phenotypes including impaired left ventricular contractility, hypertrophic cardiomyopathy, adipose hypertrophy and hyperplasia [45]. miR-27a has been reported to regulate several genes involved in adipogenesis and lipid metabolism including Retinoid X receptor alpha (RXRα), ATP-binding cassette transporter (ABCA1) also known as the cholesterol efflux regulatory protein, fatty acid synthase (FASN), sterol regulatory element-binding proteins (SREBP-1 and -2), peroxisome proliferator-activated receptor (PPAR-α and −γ), Apolipoprotein A-1, Apolipoprotein B-100 and Apolipoprotein E-3 [46]. In addition, miR-27a has been reported to play a role in the regulation of *Ldlr* [47]. Our *in vitro* and *in vivo* experiments revealed that miR-27a targets *Mvk*, *Fdft1*, *Sqle*, *Ggps1* and *Mvd* in the cholesterol biosynthesis pathway (Fig.7). Consistent with our findings, administration of miR-27a mimic in *Apoe*^−/−^ mice diminished lipid levels in both plasma and peritoneal macrophages, thereby alleviating atherosclerosis by targeting macrophage-derived lipoprotein lipase (Lpl) [48]. Of note, HITS-CLIP experiments revealed that miR-27a also targeted microsomal triglyceride transfer protein (Mttp), a crucial chaperone involved in lipoprotein production [49]. Further, global pathway analysis for miR-27a targets revealed 21 pathways including steroid biosynthesis, fatty acid metabolism and biosynthesis (Fig.S10). Thus, miR-27a contributes to global regulation of cholesterol homeostasis by targeting multiple genes in lipid synthesis, lipoprotein synthesis and uptake.

### Limitations of the study

Although our findings indicate that miR-27a mimic lowers plasma cholesterol levels considerably, there are some challenges that remain unaddressed. First, a less-/non-invasive method of delivery (viz. subcutaneous or oral route) over intravenous injections would be desirable for easy administration of the mimic. Second, improved lipid formulations or nanoparticle-mediated delivery would considerably reduce the amount of mimic required and eliminate the need for frequent injections. Additionally, further chemical modifications to this mimic may attribute to increased target specificity and confer resistance to nuclease activity. Moreover, extensive assessment of biological and pharmacological effects of miR-27a mimic in animal models are necessary to evaluate its long-term safety and efficacy as a therapeutic agent.

### Conclusions and perspectives

This study identified miR-27a as a crucial regulator of cholesterol biosynthesis pathway. Egr1 modulates miR-27a expression under basal and elevated cholesterol conditions. miR-27a directly interacts with the 3’-UTR of *Hmgcr* and represses Hmgcr protein level by translational attenuation followed by mRNA decay. miR-27a augmentation in high cholesterol diet-fed *Apoe*^−/−^ mice diminished the plasma lipid and hepatic Hmgcr levels. These findings provide novel insights on the plausible role of miR-27a in the post-transcriptional regulation of Hmgcr, thereby implicating its role in cholesterol homeostasis under pathophysiological conditions.

miRNA therapeutics is an emerging and promising avenue to treat a wide array of human diseases. A fine balance between lipid/lipoprotein synthesis and lipoprotein uptake is essential for cholesterol homeostasis and dysregulation of one or more pathways could lead to drastic effects. Therefore, miRNAs targeting multiple related pathways may be more efficacious in lowering plasma lipids than the conventional approach of targeting individual proteins/pathways. In view of the ability of miR-27a to target several crucial genes in the cholesterol biosynthesis pathway, miR-27a mimic may emerge as a promising therapeutic intervention to lower plasma cholesterol. Thus, our findings and other studies provide strong impetus for further evaluation of miR-27a as a novel lipid-lowering agent.

## Abbreviations

Hmgcr: 3-Hydroxy-3-Methyl Glutaryl-Coenzyme A reductase
QTL: quantitative trait loci
MCD: methyl-β-cyclodextrin
ChIP: chromatin immunoprecipitation
RIP: ribonucleoprotein immunoprecipitation
qPCR: quantitative real-time PCR
RIPA: radio-immunoprecipitation assay
DMEM: Dulbecco’s Modified Eagle’s Medium
SREBP-1: Sterol-regulatory element-binding protein 1
PVDF: polyvinylidene difluoride
ERAD: ER-associated protein degradation
SCAP: SREBP cleavage activated protein

## ACKNOWLEDGEMENTS

This work was supported in part by a grant from the Council of Scientific and Industrial Research (CSIR), Government of India to NRM (project number: 37(1564)/12-EMR-II). This work was also partly supported by an Exploratory Research Project grant from Industrial Consultancy & Sponsored Research, IIT Madras. Research fellowships were received from Ministry of Human Resource Development (to AAK and VG), Department of Science and Technology (to VA), Indian Council of Medical Research (to SSR and HA), CSIR (to AK), Government of India. The authors are grateful to Dr. Dona Lee Wong (Harvard Medical School, Boston) for providing the Egr1 expression plasmid, and to Dr. Weihua Xiao (University of Science and Technology of China, Hefei) for the Egr1-shRNA plasmid. The authors also thank Dr. Rakesh K. Tyagi, Jawaharlal Nehru University, New Delhi, India, for providing the AML12 cell line and Dr. Madhu Dikshit, Translational Health Science and Technology Institute, Faridabad, India for help at the initial phase of this study.

## CONFLICT OF INTEREST

The authors declare that they have no conflict of interest.

## Notes

#### Summary of Updates

Manuscript file updated Supplemental files updated

