## supplementary information for "MicroRNA-27a is a key modulator of cholesterol biosynthesis"

**This PDF file includes:**

Figures S1 to S10

Tables S1 to S4

References

### SUPPLEMENTARY FIGURES

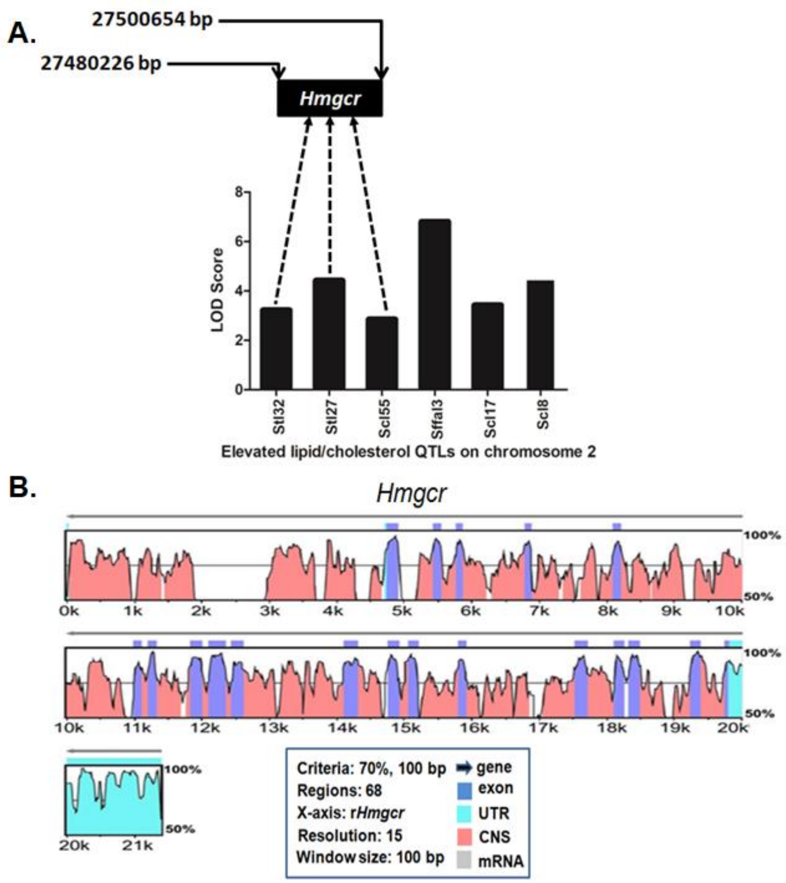


**Fig.S1. Graphical representation of rat QTLs contributing to elevated lipid/cholesterol levels and homology between mouse- and rat-*Hmgcr* gene sequences.** (A)The lipid/cholesterol-QTLs and their respective LOD scores (retrieved from Rat Genome Database) were plotted. Three of these six QTLs harbor the *Hmgcr* gene. The genomic position of rat *Hmgcr* gene is indicated. (B) Conservation analysis of rat and mouse *Hmgcr* sequences was performed using mVISTA. The horizontal axis represents the rat *Hmgcr* gene (chr2: 27480226- 27500654) as the reference sequence, whereas the vertical axis indicates the percentage homology between rat and mouse *Hmgcr* gene (chr13: 96,650,579-96,666,685). Here, window size (length of comparison) was set to 100 bp with a minimum of 70% match. Both the mouse and rat *Hmgcr* genes comprise of twenty exons; 5’-UTRs are not visible in the homology plot due to their very small sizes. CNS: conserved non-coding sequences; UTR: untranslated region.

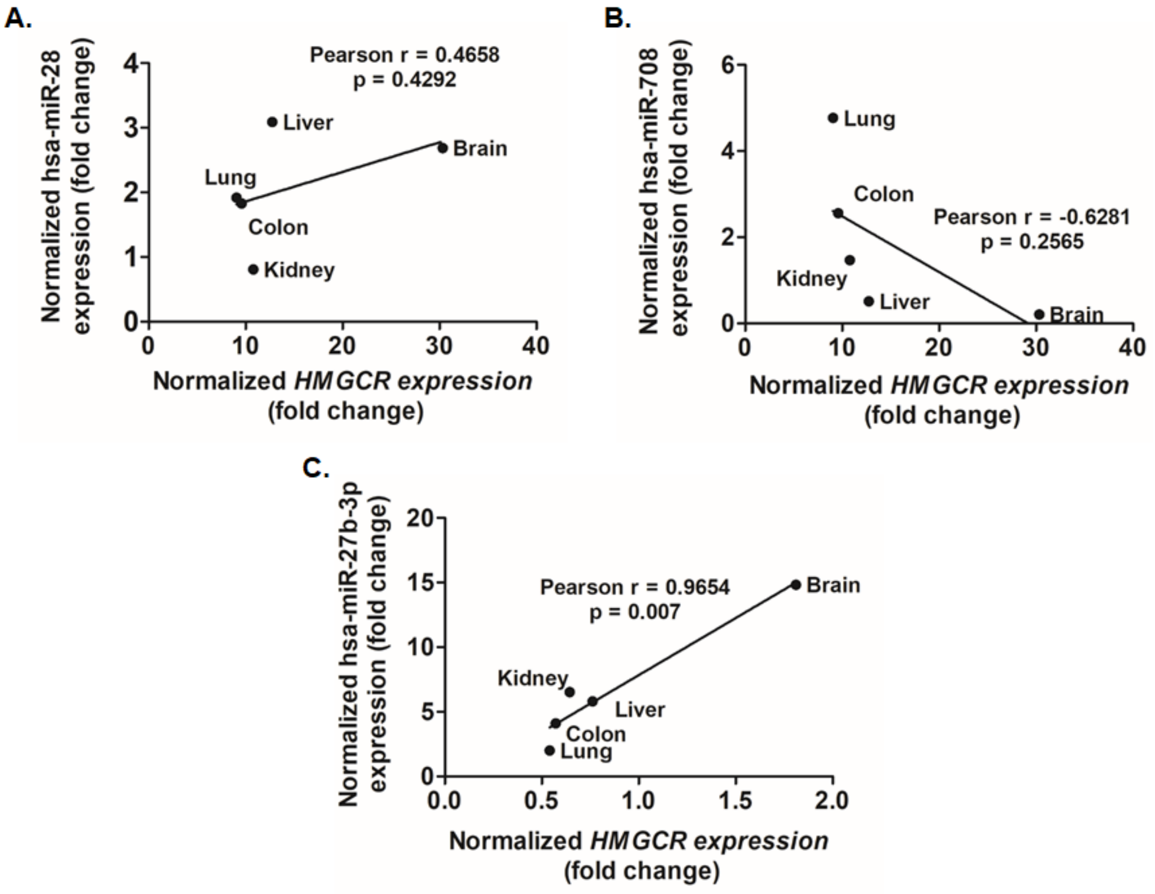


**Fig.S2: Expression analysis of hsa-miR-28, hsa-miR-708, hsa-miR-27b-3p and *HMGCR* levels in human tissues.** Endogenous *HMGCR* expression profiles in different tissues was obtained from the GTEx portal while (A) hsa-miR-28-3p, (B) hsa-miR-708 and (C) hsa-miR-27b-3p expression data was retrieved from DASHR as detailed in the materials and methods. The human Pearson r and p values for each database are shown.

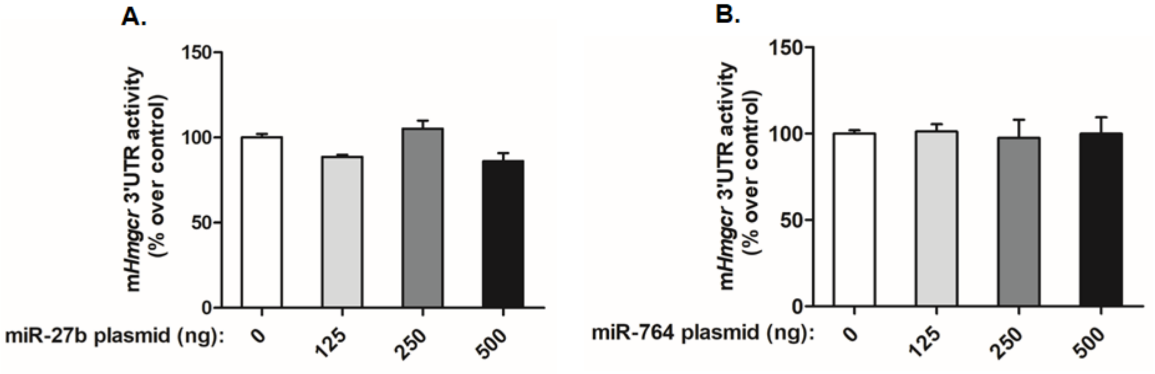


**Fig.S3. Effect of miR-27b/miR-764 over-expression on m*Hmgcr*-3'UTR.** The m*Hmgcr* 3'UTR reporter construct (500 ng) was co-transfected with increasing doses of either (A) miR-27b or (B) miR-764 expression plasmid in AML12 cells and luciferase activity was assayed. Values were normalized to total protein and are mean ± SEM of triplicate values. Statistical significance was determined by one-way ANOVA with Newman-Keuls multiple comparison test. No significant difference in the m*Hmgcr* 3'UTR activity was observed between different doses of either miR-27b or miR-764 expression plasmid.

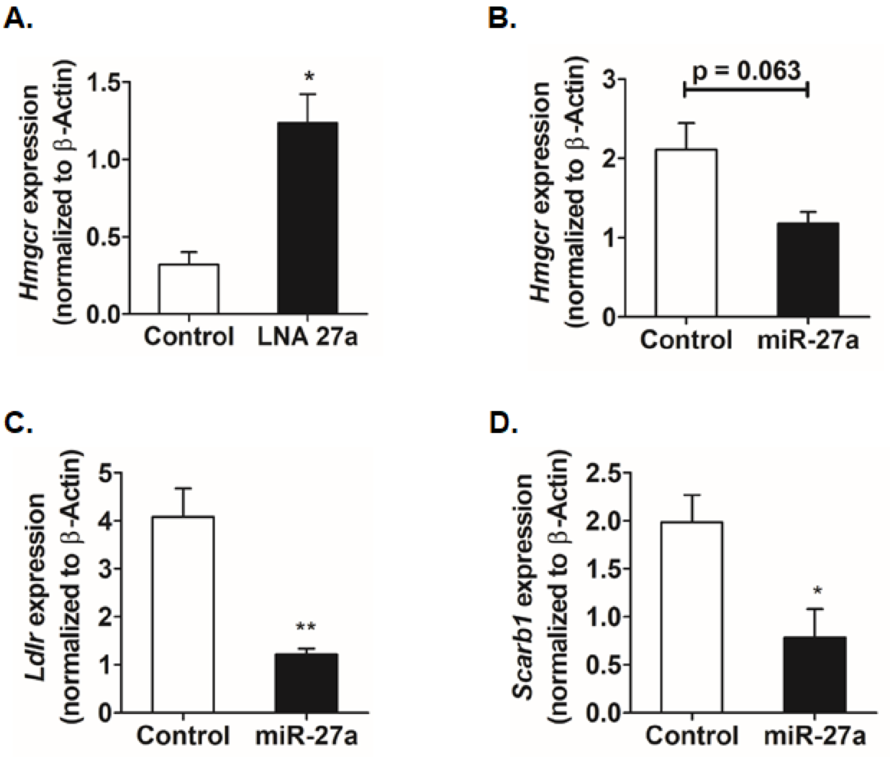


**Fig.S4. Effect of miR-27a modulation on transcript levels of Hmgcr, Ldlr and Scarb1 in cultured AML12 cells.** The relative expressions of *Hmgcr* upon transfection of (A) 60 nM of control oligo or locked nucleic acid inhibitor of miR-27a (LNA27a) or (B) 1 µg of control oligo or miR-27a mimic in AML12 cells were determined by qPCR using gene-specific primers (Table S3; n=3). The relative expression of (C) low density lipoprotein receptor (*Ldlr*) and (D) Scavenger Receptor Class B Member1 (*Scarb1*) upon transfection of miR- 27a mimic in AML12 cells was determined by qPCR using gene-specific primers. *Hmgcr*, *Ldlr* and *Scarb1* expression was normalized to β-actin mRNA in the same sample (n=3). Statistical significance was determined by Student's *t*-test (unpaired, 2-tailed). *$p<0.05$, **$p<0.01$ as compared to control.

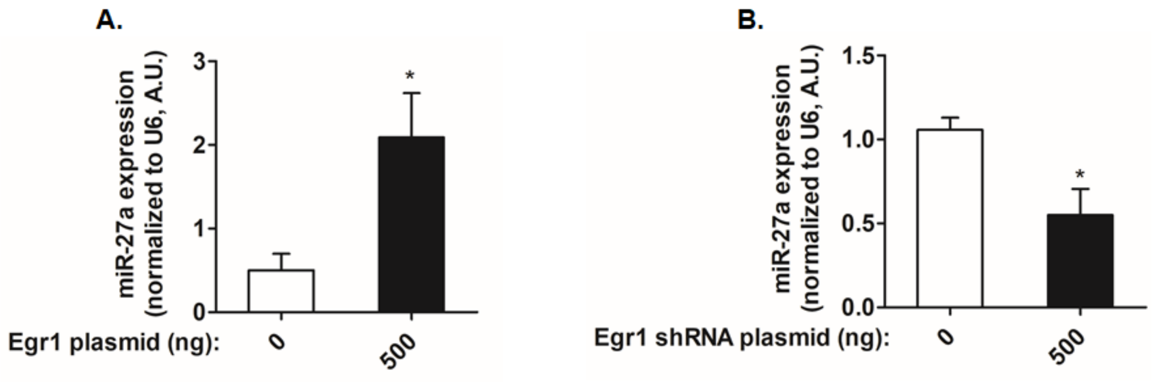


**Fig.S5. Egr1 modulates endogenous miR-27a expression in cultured hepatocytes.** AML12 cells were co-transfected with mmu-miR-27a promoter (500 ng) and 500 ng of either (A) Egr1 expression plasmid or (B) Egr1 shRNA expression plasmid followed by qPCR to probe for the endogenous levels of miR-27a (normalized to U6 RNA; n=3). Statistical significance was determined by Student's *t*-test (unpaired, 2-tailed). *$p<0.05$ as compared to the control condition.

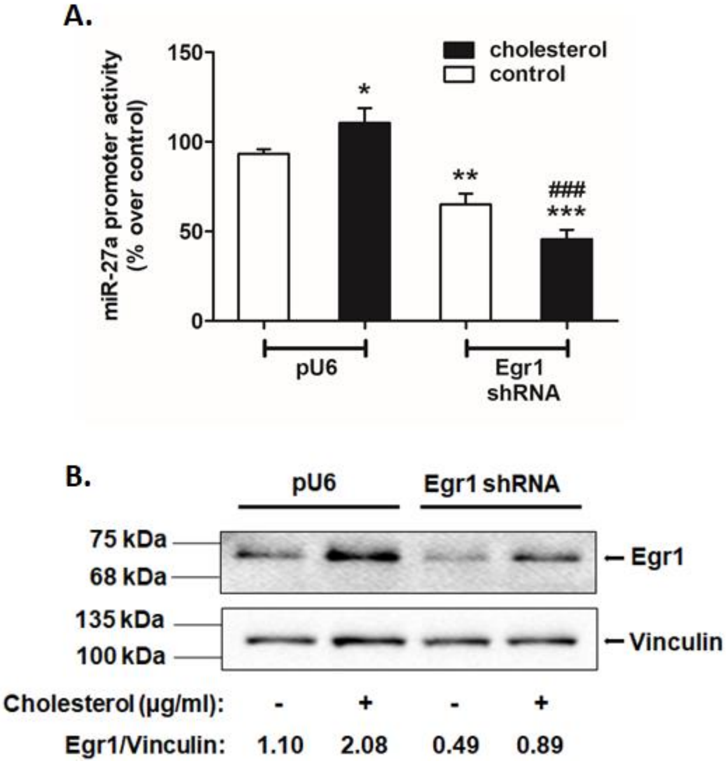


**Fig.S6. Cholesterol activates miR-27a expression via Egr1.** (A) AML12 cells were co-transfected with either pU6 (empty vector as a control) or Egr1 shRNA expression plasmid and miR-27a promoter construct (500 ng) (n=6). After 24 hrs of transfection, the cells were treated with cholesterol (20 µg/ml) for 6 hrs and lysed for luciferase and Bradford assays. The luciferase activity was normalized to total protein and expressed as % over pU6 control condition. The results are expressed as mean ± SEM. Statistical significance was determined by one-way ANOVA with Newman-Keuls multiple comparison test. *$p<0.05$, **$p<0.01$, ***$p<0.001$ as compared to pU6 control condition; $^{\#\#\#}p<0.001$ as compared to pU6 cholesterol condition. (B) Western blot analysis to probe for Egr1 levels upon transfection of AML12 cells with pU6/Egr1 shRNA plasmid followed by cholesterol treatment. The Egr1 protein levels were normalized to vinculin and indicated below the representative Western blot image. The blot shown is a representative of three experiments.

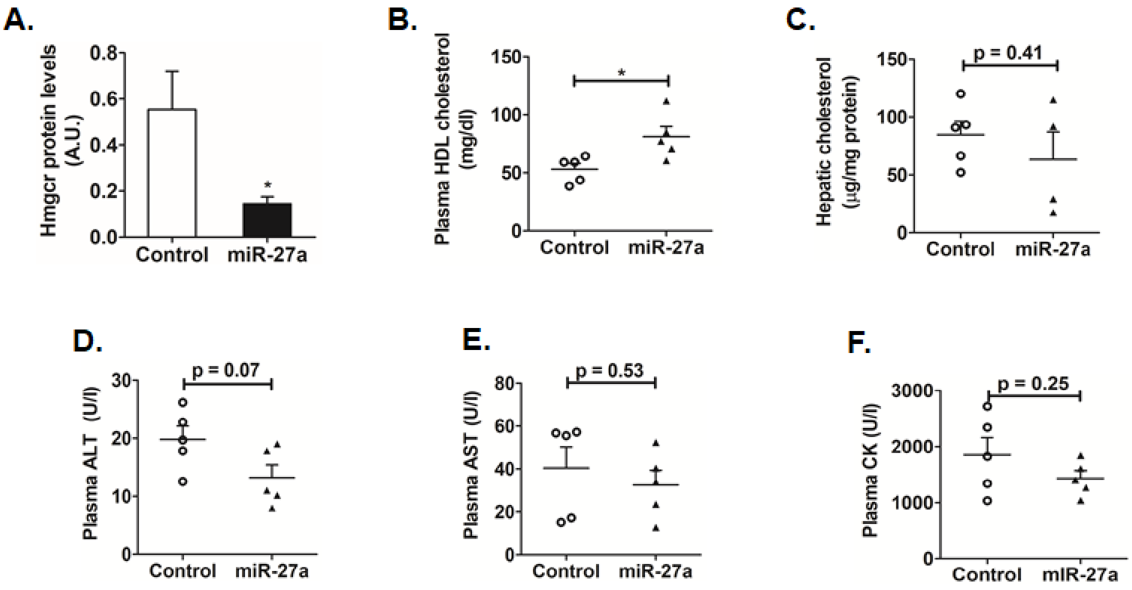


**Fig.S7. Hepatic Hmgcr, cholesterol levels and plasma biochemical parameters in miR-27a mimic and control oligo-injected *Apoe*$^{-/-}$ mice.** Total protein was isolated from the liver tissues of the aforementioned animals and was probed for Hmgcr and vinculin by Western blot analysis. (A) Hmgcr protein levels were normalized to vinculin levels and are shown in the bar plot. The blot shown is a representative of three experiments. Plasma from overnight-fasted animals was collected prior to sacrificing the animals to measure the plasma (B) HDL cholesterol (C) Lipids were extracted from liver tissue and hepatic cholesterol levels were measured in control oligo and miR-27a-injected *Apoe*$^{-/-}$ mice. The plasma (D) ALT, (E) AST and (F) CK levels were also measured. Mean ± SEM, n=5-6 animals per group. Statistical significance was determined by Student's *t*-test (unpaired, 2-tailed). * p<0.05 as compared to the control oligo-treated group.

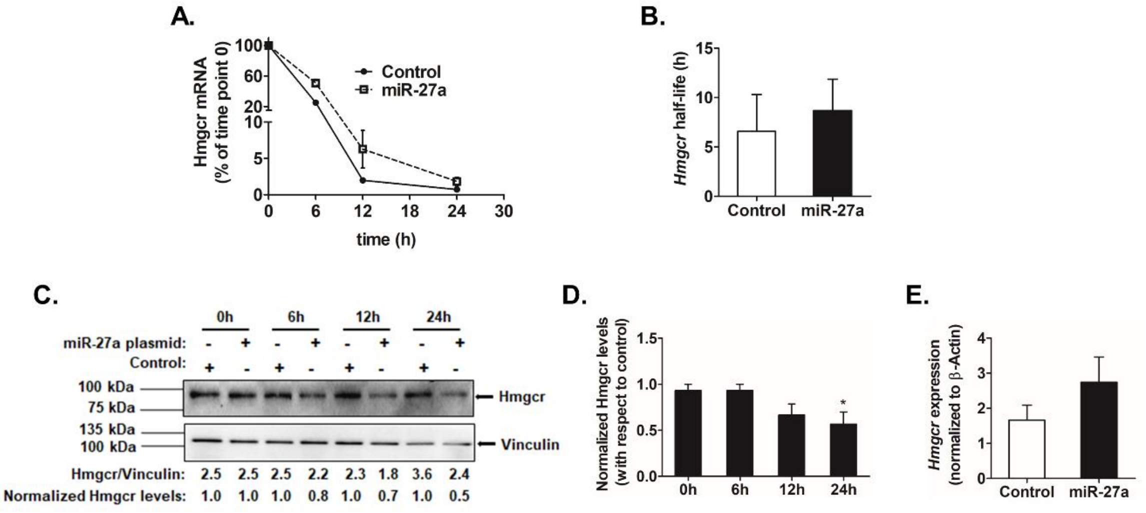


**Fig.S8. miR-27a regulates *Hmgcr* expression by translational repression in cultured hepatocytes.** AML12 cells were transfected with 500 ng of miR-27a expression plasmid or pcDNA3.1 (as control). After 12 hrs of transfection, they were incubated with actinomycin D (5 µg/ml) for different time points. (A) Hmgcr mRNA levels were plotted relative to 0 hr time point as described in Materials and Methods section (n=3). (B) Endogenous Hmgcr mRNA half-life estimation in AML12 cells on ectopic over-expression of miR-27a. The mRNA half-life of *Hmgcr* was measured over 24 hrs in the presence of 5 µg/ml of actinomycin D in control cells (transfected with pcDNA3.1) and miR-27a transfected AML12 cells. Hmgcr mRNA half-life is represented as mean ± SEM of three independent experiments. (C and D) Effect of transcriptional attenuation on endogenous Hmgcr protein levels in miR-27a over-expressed hepatocytes. AML12 cells transfected with either pcDNA 3.1 or miR-27a expression plasmid and incubated with actinomycin D for different time points. (C) Western blot analysis of total proteins was carried out probing for Hmgcr and vinculin. The relative Hmgcr protein levels normalized to vinculin for different time points are also shown. The normalized Hmgcr levels as fold change over the control for every time point of actinomycin D treatment are also indicated (n=3). (D) Bar plot showing the normalized Hmgcr protein levels expressed as fold change over corresponding control for different time points of actinomycin D treatment from three experiments. (E) The relative expression of *Hmgcr* 24 hours post-transfection of miR-27a expression plasmid in AML12 cells was determined by qPCR using gene-specific primers (n=3). *Hmgcr* expression was normalized to β-actin mRNA in the same sample. Statistical significance was determined by Student's *t*-test (unpaired, 2-tailed). *p<0.05 as compared to control.

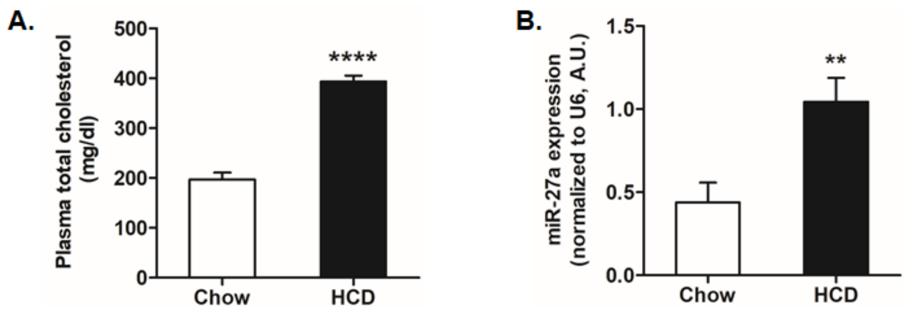


**Fig.S9. Plasma cholesterol and hepatic miR-27a levels in *Apoe*$^{-/-}$ mice fed with a normal chow/high cholesterol diet.** Male *Apoe*$^{-/-}$ mice (8-10 weeks old) were fed a normal chow diet or high cholesterol diet (HCD) regimen for 10 weeks then euthanized. Plasma levels of (A) total cholesterol from overnight-fasted animals on a normal chow/high cholesterol diet regimen. (B) qPCR for miR-27a levels in liver tissues of *Apoe*$^{-/-}$ mice fed with a normal chow/high cholesterol diet (HCD). The miR-27a expression was normalized to U6 RNA levels. Mean ± SEM, n=5-6 animals per group. Statistical significance was determined by Student's *t*-test (unpaired, 2-tailed). **p<0.01 and ****p<0.0001 as compared to control group.

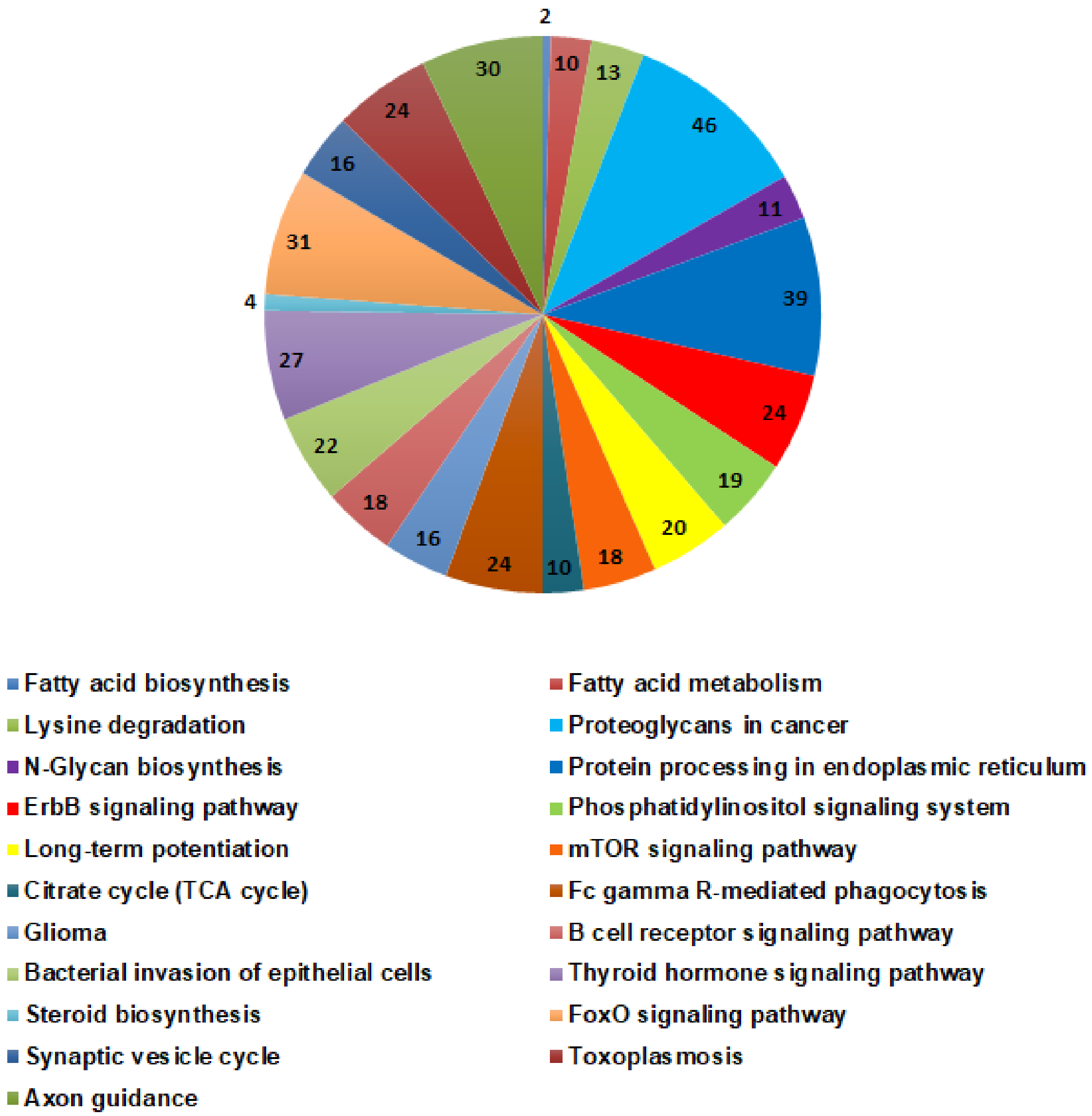


**Fig.S10. Pathways targeted by miR-27a.** Pathway analysis for miR-27a targets was performed using TargetScan (*in silico* predictions) and mirPath v3. (validated interactions). 21 pathways were commonly enriched in both these tools including steroid biosynthesis, fatty acid metabolism and biosynthesis. This analysis further demonstrates that miR-27a regulates lipid metabolism. The number of genes targeted by miR-27a in each of these pathways is indicated.

### SUPPLEMENTARY TABLES

**Table S1: Genomic position and LOD score of various lipid/cholesterol QTLs present on rat chromosome 2 (26000000- 28000000 bp) ***

| lipid/cholesterol QTLs | Start position (bp) | Stop position (bp) | LOD Score |
|---|---|---|---|
| Stl32 | 22612952 | 67612952 | 3.2 |
| Stl27 | 23837491 | 149614623 | 4.4 |
| Scl55 | 26186097 | 142053534 | 2.83 |
| Sffal3 | 27760301 | 72760301 | 6.78 |
| Scl17 | 228712271 | 266435125 | 3.4 |
| Scl8 | 231621666 | 266435125 | 4.4 |

*This data was retrieved from Rat genome database (http://rgd.mcw.edu/rgdweb/search/qtls.html?100).

**Table S2:** ***In silico*** **tools and databases employed in the study**

| Tool/database | Link | Reference |
|---|---|---|
| Rat Genome Database | http://rgd.mcw.edu/rgdweb/search/qtls.html?100 | [1] |
| VISTA | http://genome.lbl.gov/vista/mvista/submit.shtml | [2] |
| miRWalk | http://zmf.umm.uni-heidelberg.de/apps/zmf/mirwalk2/ | [3] |
| miRanda | http://www.microrna.org/ | [4] |
| TargetScan | http://www.targetscan.org/vert_72/ | [5] |
| PITA | https://genie.weizmann.ac.il/pubs/mir07/mir07_prediction.html | [6] |
| RNA22 | https://cm.jefferson.edu/rna22/Interactive/ | [7] |
| RNAhybrid | https://bibiserv2.cebitec.uni-bielefeld.de/rnahybrid | [8] |
| GTEx portal | https://www.gtexportal.org/home/ | [9] |
| miRmine | http://guanlab.ccmb.med.umich.edu/mirmine/ | [10] |
| DASHR | http://www.lisanwanglab.org/DASHR/smdb.php | [11] |
| Primer 3 | http://primer3.ut.ee/ | [12] |
| LASAGNA | http://biogrid-lasagna.engr.uconn.edu/lasagna_search/ | [13] |
| JASPAR | http://jaspar.genereg.net/ | [14] |
| mirPath v3. | http://snf-515788.vm.okeanos.grnet.gr/ | [15] |
| BioGPS | http://biogps.org/#goto=welcome | [16] |
| SAGE | https://cgap.nci.nih.gov/SAGE | [17] |
| TarBase | http://carolina.imis.athena-innovation.gr/diana_tools/web/index.php?r=tarbasev8%2Findex/ | [18] |

**Table S3: Primers used for qPCR analyses**

| Name of the primer | Sequence (5' to 3') |
|---|---|
| miR-27a-SL-primer | GTCGTATCCAGTGCAGGGTCCGAGGTATTCGCACTGGATACGACGCGGAAC |
| miR-27b-SL-primer | GTCGTATCCAGTGCAGGGTCCGAGGTATTCGCACTGGATACGACGCAGAAC |
| U6-SL-primer | GTCGTATCCAGTGCAGGGTCCGAGGTATTCGCACTGGATACGACTATGGAAC |
| mmu-miR-27-FP | ACACT TTCACAGTGGCTAA |
| U6-FP | CTGCGCAAGGATGACACGCA |
| universal RP | GTGCAGGGTCCGAGGT |
| m*Hmgcr*-FP | GAGAATGCAGAGAAAGGTG |
| m*Hmgcr*-RP | GGCGAATAGACACACCAC |
| mβ-Actin-FP | CTTCTTTGCAGCTCCTTCGTT |
| mβ-Actin-RP | TTCTGACCCATTCCCACCA |
| r*Hmgcr*-FP | ACCTGCTGCCATAAACTGGAT |
| r*Hmgcr*-RP | ACCACCTTGGCTGGAATGAC |
| rβ-Actin-FP | GCTGTGCTATGTTGCCCTAG |
| rβ-Actin-RP | CGCTCATTGCCGATAGTG |
| h*Hmgcr*-FP | TCGGTGGCCTCTAGTGAGAT |
| h*Hmgcr*-RP | TGTCCCCACTATGACTTCCC |
| m*Mvk*-FP | GGTGTGGTCGGAACTTCCC |
| m*Mvk*-RP | CCTTGAGCGGGTTGGAGAC |
| m*Fdft1*-FP | ATGGAGTTCGTCAAGTGTCTAGG |
| m*Fdft1*-RP | CGTGCCGTATGTCCCCATC |
| m*Hmgcs1*-FP | TTGAGGAGTCTGGGAATACAG |
| m*Hmgcs1*-RP | CATATCGTCCATCCCAAGAG |
| m*Ggps1*-FP | GTCATCTCCAGCAGTTCCTTC |
| m*Ggps1*-RP | TCATCTCGTCCAGCATCTTC |
| m*Scarb1*-FP | CAGGTGCTCAAGAATGTCC |
| m*Scarb1*-RP | TTTGTCTGAACTCCCTGTAG |
| m*Mvd*-FP | ATGGCCTCAGAAAAGCCTCAG |
| m*Mvd*-RP | TGGTCGTTTTTAGCTGGTCCT |

Abbreviations: FP, forward primer; RP, Reverse primer; SL, stem-loop primer

**Table S4: A list of predicted miRNAs having potential binding sites in the 3'-UTR of m*Hmgcr****

| miRNA | Seed sequence (No. of bases) | ΔΔG (PITA) | RNAhybrid ΔG (kcal/mol) |
|---|---|---|---|
| mmu-miR-124 | 7 | -10.62 | -29.5 |
| mmu-miR-28 | 8 | -10.28 | -22.2 |
| mmu-miR-345-5p | 8 | -11.08 | -32 |
| mmu-miR-351 | 8 | -16.89 | -33.6 |
| mmu-miR-708 | 7 | -14.34 | -27.4 |
| mmu-miR-27a | 6 | -10.13 | -20.7 |
| mmu-miR-27b | 6 | -10.74 | -22.6 |

* miRNAs predicted by at least 5 programs that displayed a PITA ΔΔG score of less than -10 and RNAhybrid ΔG value of -20 kcal/mol were selected
